# Deep Learning-based Modeling for Preclinical Drug Safety Assessment

**DOI:** 10.1101/2024.07.20.604430

**Authors:** Guillaume Jaume, Simone de Brot, Andrew H. Song, Drew F. K. Williamson, Lukas Oldenburg, Andrew Zhang, Richard J. Chen, Javier Asin, Sohvi Blatter, Martina Dettwiler, Christine Goepfert, Llorenç Grau-Roma, Sara Soto, Stefan M. Keller, Sven Rottenberg, Jorge del-Pozo, Rowland Pettit, Long Phi Le, Faisal Mahmood

## Abstract

In drug development, assessing the toxicity of candidate compounds is crucial for successfully transitioning from preclinical research to early-stage clinical trials. Drug safety is typically assessed using animal models with a manual histopathological examination of tissue sections to characterize the dose-response relationship of the compound – a time-intensive process prone to inter-observer variability and predominantly involving tedious review of cases without abnormalities. Artificial intelligence (AI) methods in pathology hold promise to accelerate this assessment and enhance reproducibility and objectivity. Here, we introduce TRACE, a model designed for toxicologic liver histopathology assessment capable of tackling a range of diagnostic tasks across multiple scales, including situations where labeled data is limited. TRACE was trained on 15 million histopathology images extracted from 46,734 digitized tissue sections from 157 preclinical studies conducted on *Rattus norvegicus*. We show that TRACE can perform various downstream toxicology tasks spanning histopathological response assessment, lesion severity scoring, morphological retrieval, and automatic dose-response characterization. In an independent reader study, TRACE was evaluated alongside ten board-certified veterinary pathologists and achieved higher concordance with the consensus opinion than the average of the pathologists. Our study represents a substantial leap over existing computational models in toxicology by offering the first framework for accelerating and automating toxicological pathology assessment, promoting significant progress with faster, more consistent, and reliable diagnostic processes.

Live Demo: https://mahmoodlab.github.io/tox-foundation-ui/

## Introduction

The transition from preclinical research to early-stage clinical trials in drug development is a challenging phase with a high attrition rate of drug candidates^1^. A key challenge in this transition is assessing toxicity, which accounts for 82% of drug development discontinuation at the preclinical stage^2, 3^. Drug safety is evaluated in animal models through non-clinical laboratory studies^4^, which involve a comprehensive histopathological examination of standardized tissue sections to identify and document drug-induced injury in critical organs (**Fig. 1a**). Among these organs, the liver is of particular concern due to its central role in drug metabolism, especially for orally administered compounds^5–7^.

**Figure 1:**
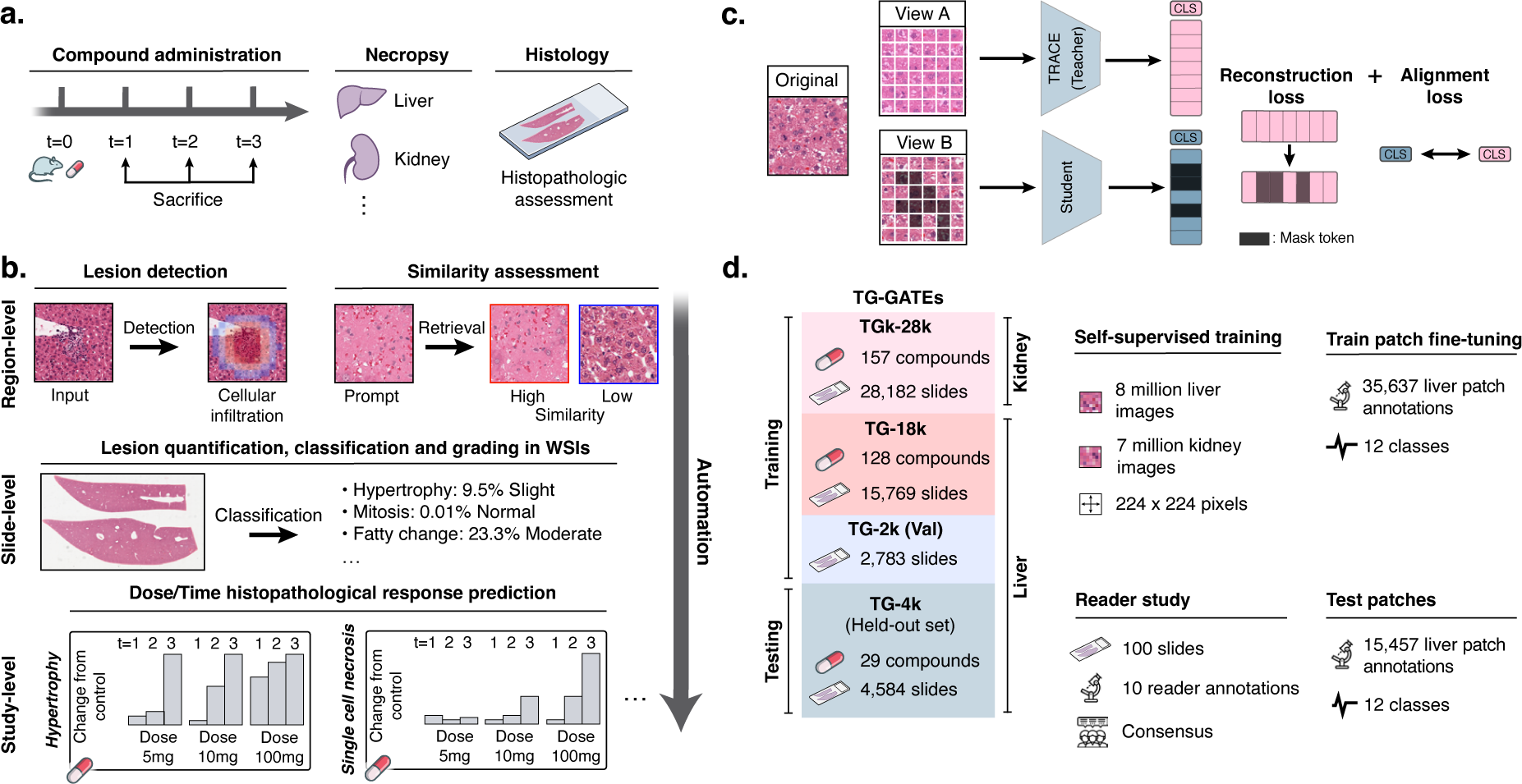
Preclinical AI-enhanced safety assessment. **a.** Before entering human clinical trials, compounds must undergo a preclinical safety assessment on rodents to assess their potential toxicity. **b.** Preclinical histopathological drug safety studies can benefit from AI assistance and automation at different scales: at *region-level* to detect and retrieve certain morphologies and lesions, at *slide-level* to automatically quantify and score abnormal lesions in WSIs, and at *studylevel* to automatically characterize the dose-time morphological response of the candidate compound. **c.** We train TRACE, a self-supervised vision encoder based on the Vision Transformer architecture trained to extract representative embeddings of small histopathological regions in *Rattus norvegicus*. TRACE uses iBOT training^32^ which combines a contrastive self-distillation objective^33^, and an image reconstruction objective^31^. **d.** TRACE is trained and evaluated on the TG-GATEs dataset. Data and annotation distribution used in this work. TG-GATEs: Toxicogenomics Project-Genomics Assisted Toxicity Evaluation System; WSI: Whole-Slide Image; AI: Artificial Intelligence; iBOT: Image BERT with Online Tokenizer.

The current approach to toxicity assessment relies heavily on manual inspection of tissue sections by pathologists, which is time-intensive and prone to high inter-observer variability^8^. This variability is particularly pronounced when diagnosing focal or ill-defined abnormalities and lesions that may be overlooked, such as focal hepatocellular necrosis, which can account for less than 1% of the tissue section. Moreover, semi-quantitative approaches frequently used in histological assessment, which involve scoring lesions on a scale (e.g., minimal, mild, moderate, severe) instead of fully quantifying them (e.g., percentage of affected tissue), further contribute to this variability ^9^. Efficient quantification requires automated tools for lesion detection, and the identification of small individual cell lesions (e.g., abnormal mitosis, single-cell necrosis) necessitates extensive review of different tissue regions at high magnification, which can quickly become prohibitive in large screening studies^10^.

These challenges contribute to the “Valley of Death” in drug development, resulting in substantial costs and unnecessary animal sacrifice^1^. Computational methods in pathology based on artificial intelligence (AI) and deep learning (DL) ^11–15^ hold promise in mitigating these challenges by accelerating safety assessment and enhancing reproducibility, as outlined by the Society of Toxicologic Pathology (STP) ^16^.

Nevertheless, existing works have primarily focused on narrow tasks, with models trained and evaluated on small cohorts ^17–26^. Considering the diversity of lesions induced by compound administration and the range of protocols used by contract research organizations, the current paradigm of developing task-specific models poses challenges to widespread adoption. The limited availability of cases for rare lesions further exacerbates this limitation. Furthermore, developing, deploying, and maintaining large ensembles of task-specific models quickly becomes impractical. This calls for the development of general-purpose and transferable models^27–30^ specifically designed for toxicologic pathology. A model trained on preclinical drug safety data could universally encode histology images, making it versatile for various tasks without the need to retrain for each task. This approach facilitates wide-ranging applicability across various studies and compounds, overall streamlining the diagnostic process.

Here, we introduce a computational framework that can address a set of diagnostic tasks for hepatotoxicity assessment across multiple biological scales (**Fig. 1b**). The bedrock of our framework is TRACE, a toxicology self-supervised model based on a Vision Transformer (**Fig. 1c**). TRACE was trained using self-supervised learning (SSL) ^31^ on 15 million image patches extracted from 46,734 whole-slide images collected from 157 preclinical studies of *Rattus norvegicus* liver and kidney tissue sections (**Fig. 1d** and **fig. S1**). TRACE is seamlessly adapted to a comprehensive set of tasks spanning histopathological response assessment, lesion severity scoring, morphological retrieval, and automatic dose-response characterization. TRACE consistently outperforms state-of-the-art baselines and clinical benchmarks from an international group of ten board-certified veterinary pathologists. Overall, TRACE introduces the first one-size-fits-all framework to accelerate and enhance toxicologic pathology, enabling significant advancements with quicker, more consistent, and more reliable diagnostic processes.

## Results

### Large-scale TRACE pretraining

TRACE is developed on TG-GATEs ^34^, a collection of histopathology slides acquired as part of the Japanese Toxicogenomics Project (JTP), a large-scale toxicogenomics research initiative. TG-GATEs consists of 157 preclinical toxicity studies of known drugs and chemicals. In each study, three doses (low, middle, high) were administered to *Rattus norvegicus* and sacrificed at different time intervals for histopathological assessment. Overall, TG-GATEs consists of 23,136 (15.1 tera-bytes, TB) liver slides (TG-23k), and 28,747 (9.9 TB) kidney slides (TG-28k) with an average of 147 slides per study (standard deviation of 38.0). In this study, we focus on liver, the principal organ responsible for the metabolism of drugs. We selected a set of 29 studies (n=4,584 WSIs) as an independent test set (TG-4k) (see **Materials and Methods**, section **TG-GATEs dataset.** and **fig. S1**). TG-4k studies were selected to encompass the various lesions and abnormalities reported by pathologists. The remaining 128 studies forms the training and validation sets (n=18,552 WSIs, TG-18k). Training and testing on different studies prevent data leakage from compound and dose-specific signatures, study-specific staining protocols, etc., all of which might lead to a lack of generalization to other studies and artificially inflate performance. From TG-18k, we randomly sampled seven million liver image patches for training our self-supervised learning model, TRACE. To increase data diversity, we additionally extracted eight million kidney image patches sampled from all 157 studies. In total, TRACE was trained on 15 million patches extracted from 47,227 different WSIs. TRACE uses Vision Transformer-Base (ViT-B) architecture trained following the iBOT training recipe ^31^ (see **Materials and Methods**, section **Vision encoder pretraining**).

### Weakly-supervised lesion classification

We investigated TRACE predictive performance on a 5-class slide-level lesion classification task. The classes were selected to encompass common lesions in rodent liver, namely necrosis, hypertrophy, increased mitosis, fatty change, and bile duct and oval cell proliferation (**table S1 and 2**, and **Materials and Methods**, section **Lesion characterization**). We follow the Multiple Instance Learning (MIL)^35, 36^ paradigm for weakly supervised slide classification. MIL consists of tessellating the slides into small non-overlapping patches, extracting patch embeddings using a pretrained vision encoder (such as TRACE), and learning an aggregation function to derive a slide embedding, subsequently used for slide classification. Here, we compare patch embeddings from TRACE with two other pretrained vision encoders, namely a ResNet50 network pretrained on ImageNet (ResNet50-IN)^36–38^, and CTransPath^39^, a Swin-Transformer-Tiny pretrained on 17 million human cancer image patches from public slide archives. We design a novel MIL model based on the attention mechanism, denoted as AttnPatchMIL, which enables deriving both slide classification along with patch-level predictions, all using slide supervision only. By following this approach, we can visualize class-wise predictions as heatmaps, which can be interpreted as segmentation predictions. A comparison with additional MIL approaches is presented in **fig. S2**, along with a description of each vision encoder and MIL strategy in the **Materials and Methods**, section **Vision encoder comparison** and **Weakly-supervised slide classifiers**.

We used TG-18k development set that was further split into train (n=15,769 WSIs) and validation (n=2,783 WSIs) set using a multilabel lesion-stratified approach based on iterative stratification for weakly supervised classification (see **fig. S1**). For testing, we used TG-4k without any form of domain adaptation or stain normalization. We evaluated and compared models using macro-AUC, balanced accuracy, and F1 score, with an additional report of 95% confidence intervals using 100 bootstrapping examples (see **Materials and Methods**, section **Evaluation Setting**). TRACE combined with our MIL method, AttnPatchMIL, reaches 96.9% multilabel macro-AUC, and shows better performance than CTransPath and ResNet50-IN. Foremost, TRACE outperforms CTransPath by 5.0% and ResNet50-IN by 8.1% as shown in **Fig. 2a**). This observation holds for other metrics in TG-4k (**Fig. 2c**). When analyzing class-wise performance, we observe that TRACE outperforms baseline encoders on all five lesions, with the largest gain observed in fatty change and mitosis detection (**Fig. 2b**). We also experimented with different MIL strategies, such as the widely employed ABMIL ^35^ and MeanMIL (constructed by taking the average of the patch embeddings), as shown in **fig. S2**. Our observations remain in that TRACE consistently outperforms other encoders using ABMIL and MeanMIL. Overall, AttnPatchMIL provides similar or slightly better performance than ABMIL irrespective of the underlying feature extractor, for instance, 95.6% *vs.* 96.9% using TRACE features (**fig. S2f,g,h**). Interestingly, and despite its simplicity, MeanMIL delivers a performance of 91.2% AUC, only 4.4% lower than ABMIL (**fig. S2i,j,k**). This illustrates the quality of TRACE features in encoding distinct and informative morphological patterns.

**Figure 2:**
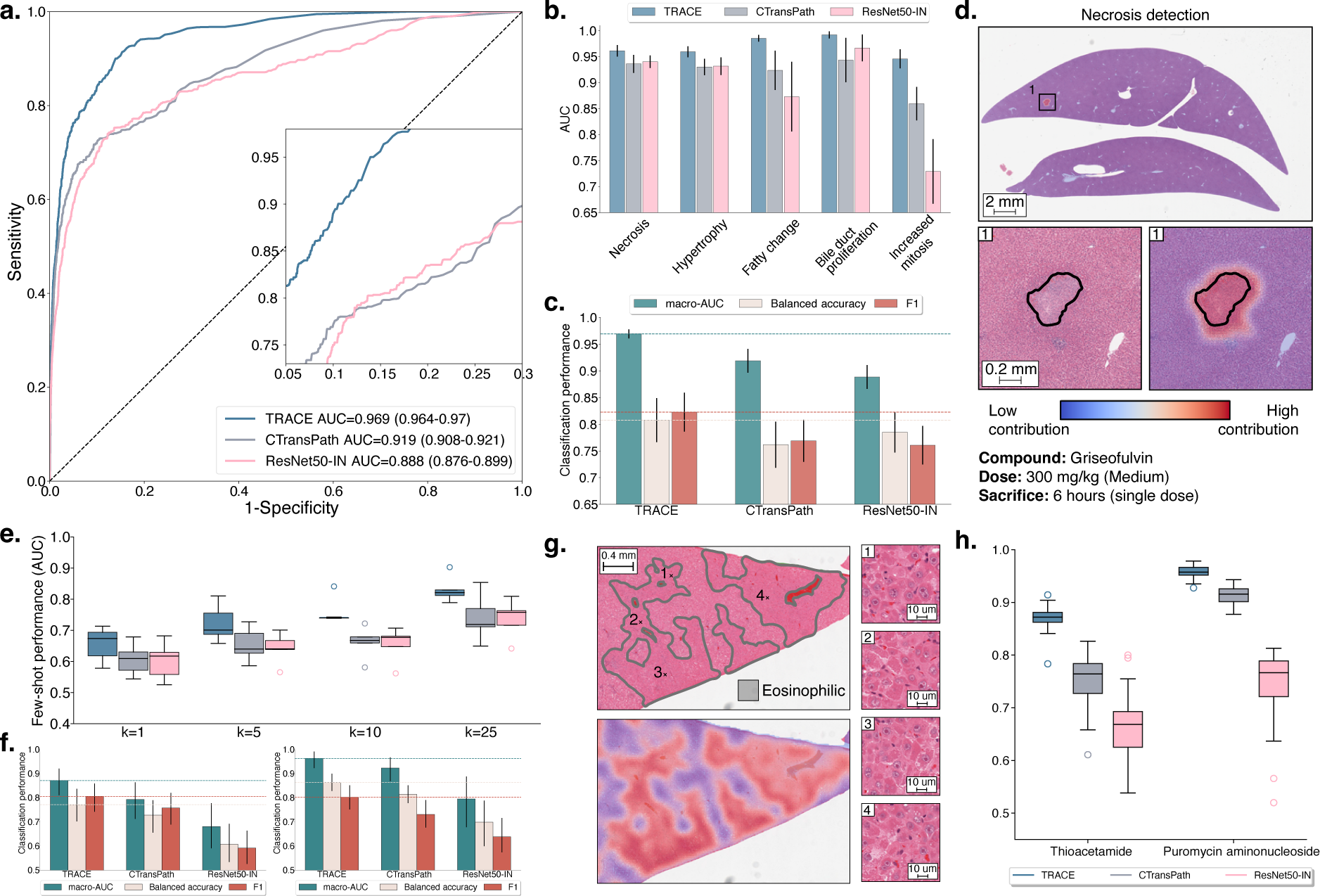
Weakly supervised slide classification. **a.** Multi-label AUROC and macro-AUC comparisons of TRACE against ResNet50-IN and CTransPath encoders, with an AttnPatchMIL model for 5-class lesion classification. **b.** Class-wise AUC of AttnPatchMIL. **c.** Macro-AUC, balanced accuracy, and F1 comparisons of TRACE against other encoders. **d.** Visualization of necrosis attribution using AttnPatchMIL and 80% patch overlap. Blue represents low contribution, and red represents high contribution. **e.** Few-shot learning performance of TRACE against ResNet50-IN and CTransPath evaluated using linear probing. Slide embeddings are defined by taking the average patch embedding (MeanMIL), and are evaluated using macro-AUC for varying numbers of training samples *k* for each class. **f.** Single-slide prompting for detecting eosinophilic changes in thioacetamide (left) and basophilic changes in puromycin aminonucleoside (right). **g.** Visualization of the similarity between the slide prototype and each patch embedding of a high-dose sample with eosinophilic cellular alteration. Results are shown as a pseudo-segmentation map indicating the presence of the morphology of interest. **h.** Single-patch prompting classification of basophilic and eosinophilic cellular alteration. Mean and standard deviation reported over ten single-patch prompts. Detailed performance metrics for all slide classification tasks are further provided in **Materials and Methods**. All models are trained on TG-18k and tested on TG-4k. Error bars in **a,b,c** represent 95% confidence intervals and were computed using non-parametric bootstrapping (100 iterations). Error bars in **e,f,h** represent standard deviation and were computed using classification performance repeated over 10 runs.

As AttnPatchMIL can derive patch-level predictions at no additional training cost, we can visualize patch attribution for each individual lesion and understand the model’s internal behavior. This property is enabled by the AttnPatchMIL aggregation function, which derives the final slide-level logits as a sum of patch-level logits (interpreted as patch predictions). Visualization of patch-wise attribution in a sample exposed to 300mg/kg of griseofulvin reveals the method’s efficiency in locating and segmenting necrotic regions (**Fig. 2d**). Additional examples in **fig. S2e** provide attribution maps for fatty change and hypertrophy in a sample exposed to a daily dose of 100mg/kg of methylene dianiline. Both lesions are clearly delineated by AttnPatchMIL, showing the versatility of the method.

### Data and label scarcity evaluation

Some compounds may induce rare morphological lesions for which only a small collection of training examples is available, thereby rendering weakly supervised classification methods inappropriate. For instance, while some lesions such as cellular necrosis (n=583, 2.5%) are common in liver, other ones are almost never encountered, such as giant cell hepatitis (n=1, 0.004%) or mineralization (n=6, 0.026%). Other lesions can be compound-specific; for instance, in TG-4k, a single compound (puromycin aminonucleoside) induces hepatocellular basophilic changes (basophilia of hepatocellular cytoplasm). Therefore, vision encoders must highlight strong fewshot learning (the ability to learn with limited training examples) and morphological retrieval capabilities.

#### Few-shot lesion classification

We designed a few-shot learning setup with *k* training examples per class. By doing so, we can evaluate the generalization capabilities and label efficiency of TRACE when only a limited number of annotations is available. Specifically, we derived slide-level embeddings for all vision encoders by taking the mean patch embedding, resulting in a fixed-dimensional and compressed representation of each slide. This representation was fed to a logistic regression classifier for training using *k ∈ {*1, 5, 10, 25*}* examples per class (a setting called linear probing). As performance may vary depending on the selected training examples, we repeated experiments over ten runs, each time sampling different examples. *K* samples were randomly drawn from TG-18k during training and tested on TG-4k for 6-class lesion classification (**table S2**)

TRACE delivers the highest performance compared to CTransPath (absolute AUC gain of +9.1% for *k*=10) and ResNet50-IN (+9.9% AUC for *k*=10) (**Fig. 2e**). The standard deviation for all vision encoders increases with smaller values of *k*, as expected from random slide selection during training, which can include small lesions and study-specific morphological features that render generalization challenging. However, as we increase *k*, the standard deviation decreases, and the overall performance increases (absolute AUC gain of +11.3% from *k*=5 to *k*=25 with TRACE). TRACE trained with *k*=1 shot (AUC 64.4%) delivers similar performance as CTransPath with *k*=10 (AUC 65.6%) and ResNet50-IN (AUC 64.9%), which demonstrates the superior label-efficiency of TRACE compared to other encoders. A similar observation holds for other evaluation metrics such as balanced accuracy and F1 score, where TRACE outperforms CTransPath by 6.4% balanced accuracy and 5.2% F1 score for *k*=10 (**fig. S3.a,b.**). When investigating the lesion-wise performance (**fig. S3c,d**), we observe that TRACE leads to the best performance across all lesions and all metrics. Still, we observe that large differences persist across lesions. For instance, bile duct proliferation, which has a distinct and consistent morphological signature, is the best detected (90.2% AUC with *k*=25). On the other hand, detecting small lesions such as single-cell necrosis is more challenging. We hypothesize that this issue is particularly prevalent when minor lesions occur alongside more noticeable ones, causing the classifier to concentrate on the more prominent lesion’s characteristics instead.

#### Morphological retrieval

In addition to few-shot learning, TRACE can be used for morphological retrieval (the ability to retrieve morphologies similar to an exemplar). To study this capability, we introduce visual prompting, where we use prototypes – a form of visual morphological descriptor representing each distinctive category – to retrieve morphological regions similar to the prompt. Visual prompting is analogous to text prompting, which aims to retrieve the most similar regions corresponding to a textual description. Here, we experiment with two types of visual prompting: single-slide (**fig. S4a**) and single-patch prompting (**fig. S4b**). In single-slide prompting, we use an entire slide as a prompt. Once the slide containing the lesion of interest (referred to as the positive slide) is identified, a slide prototype is formed by taking the average patch embedding. In single-patch prompting, we instead identify an image patch (a 256*×*256 pixels region) representing a canonical exemplar of the lesion of interest from pathologist’s annotations. For both types of prompting, we compute the cosine similarity metrics between the prompt and all patch embeddings in a test slide and average to get the similarity score (**fig. S4c**). Visual prompting obviates the need to train a complex dedicated architecture by simply relying on similarity metrics in the embedding space between the prompt and test slides. As prototyping requires a single training example (patch or slide), it can be considered a “one-shot” learning strategy. Additional information is provided in the **Materials and Methods** section **Visual prompting**.

We evaluate single-slide and single-patch prompting for detecting eosinophilic cellular alteration in thioacetamide and basophilic cellular alteration in puromycin aminonucleoside (see **Materials and Methods**, section **Lesion definitions**, **Fig. 2f,g,h** and **fig. S4d,e**). Specifically, we randomly select a single slide used for prompting from each considered study and evaluate retrieval on the remaining slides. To mitigate sampling bias, we repeat this operation ten times. TRACE delivers high predictive performance, where prompting a single positive slide yields an AUC of 96.3% (n=148 slides) for detecting basophilic changes. TRACE also outperforms CTransPath and ResNet50-IN by a large margin on both studies when evaluated using AUC, balanced accuracy, and F1 score with +7.8% and +19.0% absolute AUC gain in thioacetamide (n=159 slides) compared to CTransPath and ResNet50-IN, respectively. This underscores the promise of using prompts for fast and reliable human-in-the-loop lesion detection. Prompting also enables deriving patch similarity maps, which allow for similarity visualization and lesion quantification. For instance, in eosinophilic cellular alteration detection in thioacetamide, the patch similarity aligns almost perfectly with region annotations (**Fig. 2g** and **fig. S4d**).

Instead of prompting an entire slide, we can prompt a single patch as a prototype. To assess this approach, we repeat a similar process, where we randomly sample ten positive patches (each extracted from a different slide) that we use as a prompt for retrieval. Single patch prompting also delivers high predictive performance (AUC of 87.0% and 95.8% in thioacetamide and puromycin aminonucleoside, respectively). It also outperforms other vision encoders by a large margin with an absolute AUC gain of +11.5% and +20.7% compared to CTransPath and ResNet50-IN in thioacetamide (**Fig. 2h** and **fig. S4e**). Morphological retrieval extends other lesions and compounds, such as necrosis and fibrosis detection in monocrotaline (**fig. S5**).

Overall, visual prompting via patch or slide prototyping offers easy integration for quick and efficient lesion detection, classification, and quantification. Pathologist workload can be reduced by simplifying diagnosis to a small selection of positive slides from a study, which can be used as a prompt to diagnose the remaining slides. Slide prompting only requires slide-level labels for prototype construction, whereas patch prompting requires a small tissue region that contains the lesion of interest. When the lesion comprises only a small portion of the slide (as can be the case in necrosis or fibrosis), patch prompting enables more precise visual prompting and ensures that the prompt solely encodes the morphology of interest.

### Patch classification

Despite the promising performance of weakly supervised learning, challenges remain with detecting small lesions such as extramedullary hematopoiesis (formation and activation of blood cells outside the bone marrow), which may only cover a small area of a slide and often appear alongside various other morphological features. Such setting makes it difficult to distinguish these lesions using slide-level supervision trained with MIL. To overcome this issue, we fine-tuned TRACE with patch-level annotations collected from TG-GATEs development set to enable explicit lesion predictions at the patch level. This setup differs from AttnPatchMIL in weakly supervised learning, where patch-level predictions are implicitly generated from slide-level predictions.

To this end, we gathered a mix of publicly available and new annotations (**Materials and Methods**, section **Vision encoder fine-tuning**). In total, we obtained 29,442 patches containing at least one lesion extracted from 4,768 different slides and 13,888 normal patches extracted from 3,531 different slides. The patches containing lesions were spread across twelve lesions frequently encountered in drug safety studies (**table S2 and S3**, **Materials and Methods**, section **Lesion characterization**). These lesions display a broad spectrum of morphological variations in terms of cell type, size, and cytoplasmic and nuclear alteration. Annotations include morphological features defined at the cellular level, such as mitosis and single-cell necrosis, as well as lesions affecting groups of cells, such as cellular infiltration and bile duct proliferation. Additionally, we examine extensive lesions like hypertrophy, which can cover large portions of the slide. This diverse range allows for a thorough characterization and quantification of morphological abnormalities detected in the slide.

We fine-tuned TRACE using all patch annotations from slides extracted from TG-18k using a multilabel binary cross-entropy objective with the twelve lesions of interest. We used a class-stratified 80/20% train/validation split and then reported performance on patches extracted from TG-4k. By following this strategy, we keep a similar evaluation to that of weakly supervised classification, without any data leakage from one study to another. The resulting model is denoted as TRACE (FT). We compare it against three baselines, TRACE, CTransPath, and ResNet50-IN, followed by a set of 12 logistic regression models (linear classifiers), one for each lesion, which predict the probability of lesion existence from the patch embedding. We observe that TRACE (FT) and TRACE outperform other vision encoders in terms of AUROC and macro-AUC with 98.9% for TRACE (FT) against 96.3% for CTransPath and 86.8% for ResNet50-IN (**Fig. 3a,b**). Even without fine-tuning, TRACE leads to high performance with 98.2% AUC. A detailed breakdown of class-wise predictions of TRACE (FT) and baselines further demonstrate the superiority of in-domain fine-tuning, especially for small lesions such as single-cell necrosis and mitosis (**Fig. 3** and **fig. S6**). Challenging lesions typically occupy a tiny portion of the patch and can co-appear with other more distinct features, such as cytoplasmic alterations. The detection, therefore, necessitates a powerful vision encoder trained with the fine-grained supervisory morphological cue, which TRACE (FT) aptly fits.

**Figure 3:**
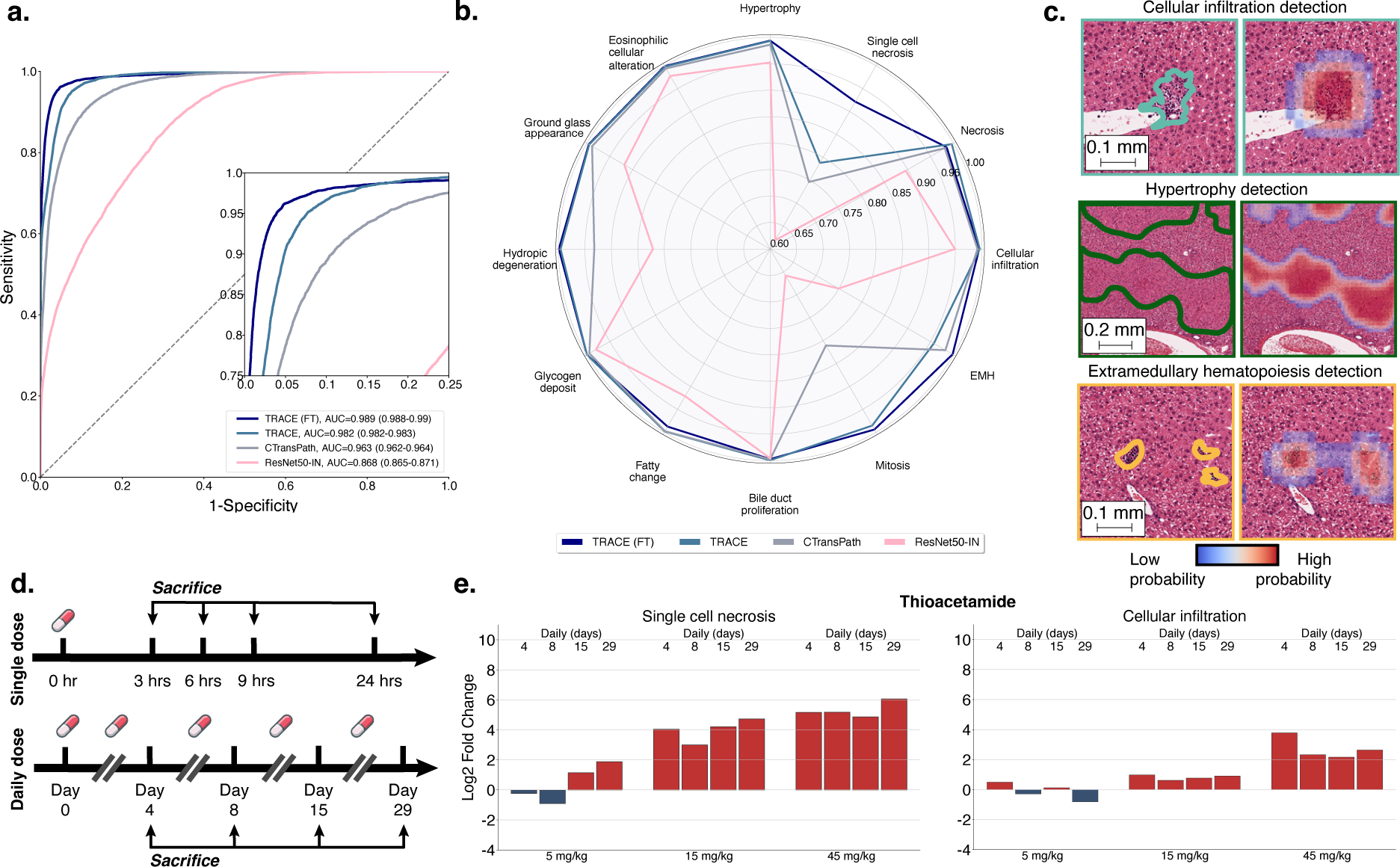
Patch-level lesion classification with TRACE fine-tuning. **a.** AUROC and macro-AUC of TRACE fine-tuned (TRACE (FT)) compared to TRACE, CTransPath and ResNet50-IN across 12 lesions-of-interest. Models are trained on patch annotations from TG-18k and evaluated on patch annotations from TG-4k. 95% confidence intervals were reported using non-parametric bootstrapping (100 iterations). **b.** Class-wise predictions of TRACE (FT) and baselines evaluated using AUC. **c.** Patch prediction visualization on a test sample from TG-4k using 80% patch overlap. Heatmap overlay highlights regions with a high probability of being a lesion, colored contours highlight lesions annotated by pathologists. **d.** Overview of time-dose toxicity assessment. Each sample group (five animals) represents the administration of a specific compound at a predetermined drug dosage level, with a predefined frequency of dosing (single or daily administration) and a distinct interval of time between administration and sacrifice. A control group is used to identify background lesions. **e.** Morphological log2 fold change of single cell necrosis (left) and cellular infiltration (right) against the control group in thioacetamide using TRACE (FT) patch predictions with 80% patch overlap.

By employing a Uniform Manifold Approximation and Projection (UMAP), we can visualize the latent space learned by TRACE (FT) across all lesioned patches from TG-4k (n=7,370) and understand the ambiguity associated with overlapping classes (**fig. S7**). This visualization underscores the task’s inherent complexity stemming from its multilabel nature due to several lesions co-occurring together (for instance, necrosis and cellular infiltration, or fatty change and glycogen deposit) and similar histomorphologic appearance (cellular infiltration and extramedullary hematopoiesis).

Furthermore, we can combine individual patch predictions to create a detailed segmentation map that reveals every lesion identified within a slide (**Fig. 3d** and **fig. S8**). To enhance the resolution of these predictions, we overlap patches by 80% and calculate the average probabilities for each patch, disregarding any probabilities below a lesion-specific threshold. This approach is particularly useful for identifying smaller lesions, like focal cellular infiltration, extramedullary hematopoiesis, or hydropic degeneration, that may constitute only a small portion of the slide.

### Automatic dose-time response assessment

Beyond single-slide diagnosis, preclinical safety assessments aim to understand and characterize the dose-response relationship of the compound. To this end, the onset of morphological abnormalities and lesions are monitored at consistent time intervals across various dosages. In TG-GATEs, five animals are used in each sample group (administration of a given dose and sacrifice time). For each identified lesion, a severity score is assigned to describe the extent of the finding. Severity score assignment is crucial to distinguish small lesions from severe drug injury. However, the reported scores generalize poorly within and across studies due to high inter-observer variability, the complexity of measuring the extent of changes, and the lack of unified standards^8^. This complicates the precise and quantifiable characterization of morphological changes, which, along with the burden of manual examination, renders drug-response assessment nontrivial even on the scale of a single study (which typically involves several hundred slides).

Instead, by leveraging the superior patch classification capacity of TRACE (FT), we can obtain quantification scores for each lesion, which we express as a percentage of the lesion area in tissue compared to its normal counterpart. Specifically, we average quantification scores of each lesion over each sample group, which enables monitoring the drug dynamics across all doses and sacrifice time points (**Fig. 3d**). As some lesions occur spontaneously, we need to normalize background lesions found in treated samples by a set of control slides. For instance, extramedullary hematopoiesis is a naturally occurring lesion in rodents, regardless of the administered compound. To this end, we compute the log2 fold change, which quantifies the relative morphological change between the control and sample group conditions on a logarithmic scale (**Fig. 3e** and **fig. S9**).

By adopting this approach, we can observe the onset of lesions as the dose and sacrifice time increase, as exemplified with thioacetamide in **Fig. 3b**, and methyltestosterone and hydroxyzine in **fig. S9**. For instance, we observe that even a single dose of methyltestosterone induces an abnormal increase in mitosis at a high dose (300 mg/kg) after 24 hours (**fig. S9a**). On the contrary, a single dose of hydroxyzine does not result in extensive lesions. However, upon administering a daily dose of 30mg/kg or more, signs of hepatocellular fatty change start developing. The severity of the fatty change escalates with an increase in dosage when administering 100mg/kg (**fig. S9b**).

Overall, using TRACE (FT) as a lesion quantification tool enables new automation possibilities for more objective and precise lesion identification and more robust drug safety assessment.

### Comparison with pathologists

To assert our model capabilities in a realistic diagnostic scenario, we designed a reader study to compare the predictions of TRACE (FT) with those of pathologists (**Fig. 4a**). To this end, a subset of n=100 slides from TG-4k (denoted as TG-100) was selected and independently reviewed by ten veterinary pathologists (average of 10.2 years of experience post-board certification). In total, slides were selected from 22 different studies (**table S4**). Slides were selected based on TG-4k original annotations to include a wide range of lesions and severity. Apart from ensuring that the selected slides did not contain any severe staining or scanning artifacts, no other selections were made.

**Figure 4:**
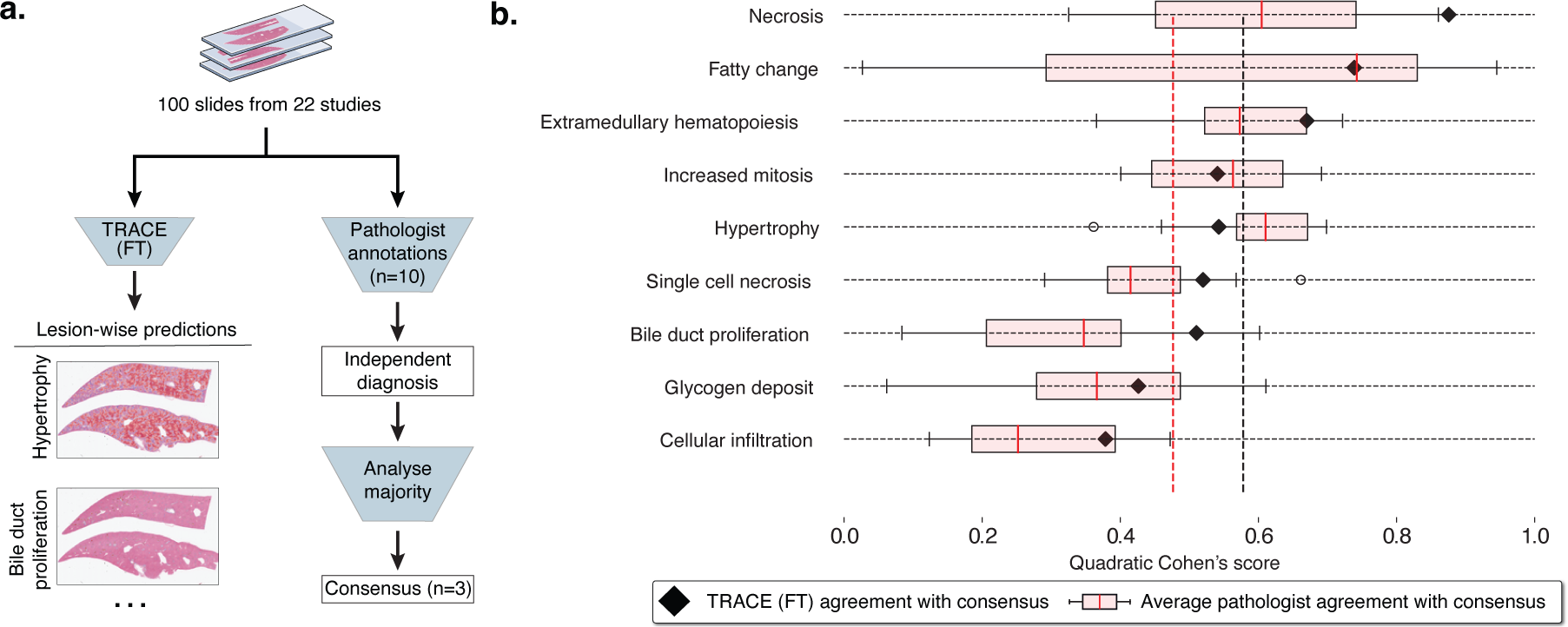
Comparison with pathologists. **a.** 100 slides from 22 studies were randomly selected from TG-4k for comparison with pathologists. Ten veterinary pathologists were asked to report the presence of each lesion and assign a score from normal, minimal, mild, moderate, or severe. After the independent evaluation, a consensus was reached between three pathologists to derive the gold standard. In parallel, each slide was processed by TRACE (FT) to derive patch predictions with 80% patch overlap. Class-wise patch predictions were then thresholded to retain solely high-confidence predictions, from which the quantitative scores describing the percentage of the slide highlighting each lesion were derived. **b.** Comparison between TRACE (FT) and consensus (black triangles), and pathologists and consensus (pink box plot). Evaluation reporting Quadratic Cohen’s kappa score. Vertical lines indicate the average TRACE (FT) and pathologist agreement with the consensus. Boxes indicate quartile values of Quadratic Cohen’s Kappa score, with the red center line indicating the 50th percentile. Whiskers extend to data points within 1.5x the interquartile range.

Based on pathologists’ expertise, nine lesions were selected for evaluation that met the following criteria: 1) lesions are relevant and frequent in toxicology studies; 2) they are reliably detectable on H&E-stained tissue sections; and 3) they are well characterized and defined by INHAND guidelines^40^ (the International Harmonization of Nomenclature and Diagnostic Criteria provided by Society of Toxicology Pathology). For instance, eosinophilic cellular alteration, ground glass appearance, and hydropic degeneration were combined with hypertrophy due to their frequent co-occurrence and unspecific toxicologic relevance as a stand-alone lesion. All pathologists were instructed to mark each lesion type by assigning it a score spanning normal, minimal, mild, moderate, and severe. After the independent annotation process, all slides were reviewed and discussed by a group of three pathologists to define a consensus. Additional information, including instructions and lesion definitions, is provided in the **Materials and Methods**, section **Reader study** and in **table S2**.

When comparing TRACE (FT) with pathologists against the consensus, we find that, on average, TRACE (FT) outperforms pathologists by +10.2%, as measured by the Quadratic Cohen’s kappa score (**Fig. 4b**, black: average TRACE (FT), red: average pathologist). The AI system performs better across 6/9 lesions. We observe large variations across lesions between pathologists and the consensus, especially for fatty change detection. Fatty change, characterized by hepatocellular vacuolation, may be similar to glycogen accumulation, which can partially explain this variability. The best-predicted lesion by TRACE (FT) is necrosis (87.6% kappa score), probably because of its distinctive and unequivocal characteristics. Even if well-characterized, the extent of this lesion might be underestimated if scored manually due to an incomplete detection when diagnosing slides rapidly. In contrast, the agreement between TRACE (FT) and the consensus for hypertrophy was reduced. This can be attributed to the ill-defined histologic appearance of this lesion, which can be affected by variations in staining, inducing ambiguity. To detect evidence of increased cell size, which defines hypertrophy, knowledge of normal cell size or the presence of unaffected tissue to compare against, ideally on the same tissue section, is required. In addition, glycogen deposit presents a lower agreement with the consensus. This was anticipated to occur for two reasons. Firstly, intracellular accumulation of glycogen is, to a certain extent, a normal finding in rodent livers as a response to food ingestion, with specific thresholds that have not been established. Therefore, it remains the pathologists’ interpretation to assess if the presence of glycogen is considered physiologic or pathologic. Second, this lesion shares histomorphologic characteristics with fatty change, as both lesions are defined by hepatocellular vacuolation. A lower agreement with consensus was also reached regarding cellular infiltration. This was considered a consequence of the histologic similarities between this lesion and extramedullary hematopoiesis, where hematopoietic cells can be difficult to distinguish from inflammatory cells. Overall, the predictive capabilities of TRACE (FT) bring new clinical tools to pathologists for more reproducible and objective assessment.

## Discussion

We introduce TRACE, a state-of-the-art Vision Transformer (ViT-Base, 86 million parameters) model to support toxicity assessment in preclinical drug safety studies. TRACE is based on the TG-GATEs dataset, the largest public cohort of histology slides of preclinical studies to date. TRACE was trained with self-supervised learning (SSL) on a dataset of 15 million image patches extracted from liver and kidney tissue sections of *Rattus Norvegicus*. In total, 47,227 whole-slide images collected from 157 preclinical studies were harnessed for training TRACE. Compared to state-of-the-art vision encoders trained on human cancer samples or natural images that are commonly used in computational pathology, TRACE consistently reaches the best performance when evaluated on several downstream tasks spanning various spatial scales, underscoring the versatility of TRACE for histopathological toxicity assessment.

On weakly supervised tasks trained with slide-level labels, TRACE largely outperforms other vision encoders and, when combined with AttnPatchMIL, highlights patch-level attribution properties that enable deriving pseudo-segmentation maps from slide-level supervision. The patch embeddings of TRACE also show remarkable retrieval capabilities, where visual prompting of a single slide or image patch enables retrieving slides sharing similar morphologies. These results demonstrate the benefits of pretraining large-scale vision encoders with large sets of in-domain images. When benchmarked against ten veterinary pathologists, TRACE fine-tuned with patch annotations compared favorably and provided better overall performance than pathologists. The reader study highlights large inter-observer variability across all lesions, which can be attributed to two main factors. First, detecting lesions on large batches of tissue sections, especially subtle, non-specific, and focal ones, is time-consuming and cumbersome. Second, quantifying lesions in slides and assigning severity scores in a consistent manner is challenging. TRACE can address these challenges with automatic detection and quantification of lesions, producing reproducible quantification scores that can generalize and transfer within and across studies.

TRACE holds the potential to revolutionize the entire histopathological diagnostic chain involved in drug safety assessment. This transformation can bring different levels of automation and assistance to pathologists, from assisting diagnosis on small *regions-of-interest* within a slide to automating the assessment of an entire *study* comprised of several hundred slides. At the level of ROIs (or image patches), TRACE can identify and outline specific lesions and morphological features of interest. By further employing annotations for individual ROIs, TRACE can identify and retrieve morphological patterns that may be present in only a few slides. When expanded to the slide level, TRACE can classify and quantify lesions. Such assistive tools can be integrated into clinical diagnosis to accelerate and enhance reproducibility from one sample to another and from one study to another. At the study level, TRACE can be employed to characterize the dose-response relationship of a candidate drug. This can be accomplished either through the absolute quantification of a set of lesions per sample group or, as traditionally performed in toxicogenomics studies^41–43^, by quantifying deviations from a control group to account for spontaneous lesions. Overall, the different levels of granularity offer varying degrees of automation, assistance, and control, thereby enabling new synergies between pathologists and TRACE. Alternatively, TRACE can serve as an independent validation mechanism, ensuring concordance between its assessments and pathologists.

Despite the wide-ranging tasks TRACE can handle, our study includes limitations. Firstly, diagnoses conducted as part of TG-GATEs were not designed for developing AI applications, leading to redundant, equivocal, and ill-defined lesion reports. While guidelines, such as IN-HAND, aim to bring a unified taxonomy of diagnoses and lesion scoring, inconsistencies remain, such as defining thresholds for reporting increases in mitotic figures or reporting nonpathological glycogen deposits, which makes the comprehensive evaluation of TRACE challenging. In addition, many lesions are very rare, with a prevalence of less than one per thousand cases, such as giant cell hepatitis, which prevents rigorous assessment of TRACE retrieval capabilities on those uncommon findings. Moreover, our study focuses on hepatotoxicity due to the liver’s primary role in drug metabolism. Nonetheless, a comprehensive safety evaluation necessitates the examination of additional organs. Additionally, our model is exclusively trained on *Rattus norvegicus*, the predominant species in preclinical evaluations^44^. However, including other species like rabbits and mini-pigs could increase the generalization of our findings.

We anticipate these challenges will be addressed as more data from preclinical drug safety studies become available, notably with joint contributions from the pharmaceutical industry and public institutions. For instance, initiatives such as the BigPicture project^45^ plan on gathering over three million slides from several pharmaceutical companies, which could be combined with the ongoing digitization effort of the National Toxicology Program (NTP), the inter-agency entity responsible for evaluating and reporting toxicology in the United States, which have gathered over 2,000 studies and 7.5 million slides. Broadly, our study lays the foundation for unlocking new avenues in AI-driven histopathological assessment of drug toxicity. Foundation models based on large-scale pretraining of diverse data offer a promising direction for assisting, augmenting, and automating diagnostic assessment. We anticipate that large-scale efforts from pharmaceuticals and academia will further accelerate these trends.

## Materials and Methods

### Large-scale visual pretraining

#### TG-GATEs dataset

The TG-GATEs dataset^34^ (Toxicogenomics Project-Genomics Assisted Toxicity Evaluation System) is a collection of histopathology and gene expression data generated by the Japanese Toxicogenomics Project (JTP) consortium as part of a large-scale toxicogenomics research initiative. This dataset was created to study the effects of various compounds (drugs and chemicals) to understand how these might cause toxicity and adverse health effects.

#### Study design

All TG-GATEs studies (157 used in this work) are based on 5-week-old male Sprague-Dawley (SD) *Rattus norvegicus* from Charles River, Japan. All SD rats were stratified into groups of 20 animals based on body weight. During the study, SD rats had free access to water and pelleted food (radiation-sterilized CRF-1; Oriental Yeast Co., Tokyo, Japan). Two types of compound administration were tested: single-dose and daily-dose administration. In single-dose experiments, SD rats were administered one dose and were sacrificed 3, 6, 9, and 24 h after administration. In daily-dose experiments, SD rats received a new fixed-dose every day for 3, 7, 14, or 28 days, and were sacrificed 24 h after final administration (4, 8, 15, or 29 days after the first dose, see **Fig. 3d**). Each unique combination of compound, dose, single or daily dose, and sacrifice was tested on five SD rats. Each compound was tested at three different doses (low, medium, and high), together with a control group. Single and daily administration experiments may be based on different doses. The ratio between low, medium, and high doses follows roughly 1:3:10. In some high-dose cases (especially in daily dose administration), death was observed before the end of the study, in which case less than five samples are available. Most drugs were selected for their known toxicity on liver, kidney, or both. In addition, some reference chemicals were included for which their toxicity is well understood. In total, 157 compounds were tested with corresponding slides and histopathological assessment.

#### Histopathology

Following sacrifice, organs were harvested for histopathological assessment. More information on the liver sampling site can be found in ^34^. After slicing, the liver and kidney sections were stained with H&E (hematoxylin and eosin) and mounted on glass slides. Slides were then scanned using an Aperio scanner at 20*×* magnification (0.49*µ*m/px). All slides were downloaded from the publicly available Open TG-GATEs portal^1^. In total, 23,136 (15.1 tera-bytes, TB) liver slides, referred to as TG-23k, and 28,747 kidney slides (9.9 TB) were downloaded with, on average, 147 slides per study (*±*38.0). The average image size is 54852 *×* 33564 pixels at 20*×* magnification (or 26.9 *×* 16.4 mm sections).

#### Lesion characterization

TG-GATEs histopathology data are accompanied by annotations describing the lesions identified in slides. In total, 66 different liver lesions were identified across 23,136 WSIs, many of which are synonyms or sub-categories of lesions, such as “Degeneration, fatty” or “Vacuolization, cytoplasmic” to describe a fatty change in hepatocytes. The lesion distribution is highly skewed with common ones, such as necrosis (n=583, 2.5%), and very rare ones, such as mineralization (n=6, 0.026%). We organized lesions into different sets used for evaluating TRACE under various scenarios as described in **table S2**.

The scope and definition of each lesion used in this study are based on the INHAND guidelines^40^ (International Harmonization of Nomenclature and Diagnostic Criteria). Four main categories were studied, with one to six different lesions in each category. The majority of lesions are linked to hepatocellular responses, cellular degeneration, injury, and death, followed by the group of non-neoplastic proliferative lesions. Half of the lesions have a characteristic histomorphology and thus are well-defined as such, including necrosis, cellular infiltration, bile duct proliferation, increased mitosis, and extramedullary hematopoiesis. Less specific histologic appearance is typical for the remaining six lesions, which are primarily defined by hepatocellular cytoplasmic alterations. In fatty change, hydropic degeneration, and glycogen deposit, vacuolation or rarefaction of the cytoplasm is a key feature. The distinction of these three lesions may be difficult based on H&E-stained tissue sections, and interobserver variability is therefore expected to occur. Other less specific lesions include basophilic and eosinophilic cellular alteration and ground glass appearance, which are defined by cytoplasmic tinctorial and texture changes. Hence, the detection of these lesions relies very much on the tissue and staining quality, which makes them prone to inconsistent reporting by pathologists. A summary of lesions considered in this study is provided in **table S2**.

#### Slide split

To mitigate batch effects, we split training and testing data at study level, such that two slides from the same study (same compound administered) cannot be found in different splits. We selected 29 studies (n=4,584 WSIs) encompassing the variability of lesions, which we denote as TG-4k. The remaining 128 studies (n=18,552 WSIs) are used for training and validation, which we denote as TG-18k. The TG-18k is further split into train (n=15,769 WSIs) and validation (n=2,783 WSIs) set using a multilabel lesion-stratified approach based on iterative stratification (see **figure S1**).

#### Vision encoder pretraining

TRACE was trained on 15 million patches extracted from 46,734 WSIs, including 28,182 kidney and 18,552 liver slides, accounting for a total of 8,144,093 kidney image patches and 6,917,697 liver image patches. Patches were randomly sampled without any form of patch selection.

At the core of our AI-enhanced drug safety pathology assessment is a large-scale self-supervised learning (SSL) model based on a Vision Transformer (ViT). ViTs have shown remarkable representation learning abilities in vision by extending the principles of Transformers ^46^, originally developed for natural language processing, to images ^47^. ViTs use self-attention to capture spatial relationships among small regions (or tokens) of the input. Specifically, we train a ViT to compress image patches of 256*×*256 pixels into 768-dimensional representations that can further be used for downstream tasks, such as image and slide classification or retrieval. In this work, we train a ViT using iBOT^31^, a state-of-the-art training strategy based on student-teacher knowledge distillation. iBOT combines two objectives: First, a self-distillation objective that aims to align different representations (or views) of the input image. This objective enables capturing high-level context and semantics information of the image (for instance, to build stain-invariant or rotation-invariant representations). Secondly, a masked image modeling (MIM) objective that aims to reconstruct small regions (or tokens) from the neighboring regions. This objective enables capturing the image’s internal structure, similar to masked language modeling in Large Language Models ^48^ and has successfully been adapted for images ^49^.

Formally, we build two representations (or views) of an input image *x*, denoted as *u* and *v*, using augmentation techniques, such as color jittering, mirroring, and zooming. Each view is then randomly masked using blockwise masking^49^, where a series of blocks (rectangles of 16 patches) with a random aspect ratio are generated until a pre-defined percentage of the input is masked. We denote the masked augmented views as *u*^ and *v*^, respectively. The two views *u*^ and *v*^ are further tokenized into a set of small tokens of 16*×*16 pixels and are passed as a sequence through a teacher network (a ViT). Likewise, the tokenized sets for the two masked views *u*^ and *v*^ are passed through a student network (another ViT). The self-distillation objective (originally proposed in DINO^33^) is then defined by computing the cross-entropy loss between the [CLS] token (the image-level representation) from the teacher network and the [CLS] token from the student network. This can be seen as aligning high-level representations of masked and unmasked views. The masked image modeling objective uses learnable projections of the masked tokens from the student network to predict projections of the patch tokens from the teacher network. The projected patch tokens from the teacher network can be seen as a visual tokenizer for each masked token. The teacher network is then updated using an exponential moving average from the student network. The trained teacher network corresponds to TRACE.

During SSL pretraining, assessing the quality of the latent space being learned and knowing when to stop training can be challenging. This is particularly true when using complex loss functions composed of multiple objectives. Previous works have relied on monitoring the downstream task performance at regular intervals, for instance, after each epoch. However, such evaluation is computationally expensive, with a risk of overfitting to the downstream task. Instead, we use the rank of a representative set of 10,000 patch embeddings sampled from the test set as a quick measure to assess the embedding space quality and use it to guide hyperparameter tuning^50^. Intuitively, a higher rank signifies more diverse patch embeddings and ensures that representations have not collapsed to a few modes, *i.e.,* low rank. Here, we compute the rank as the entropy of the *d* (assuming *d < n*) L1-normalized singular values of the slide embedding matrix *H ∈ R^n×d^*, which can be expressed as:

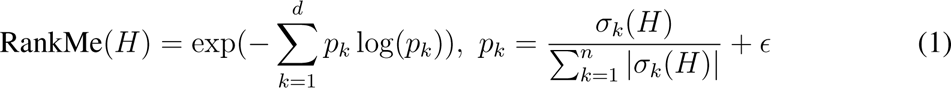

where *σ_k_* denotes the *k−*th singular of *H* (sorted from large to low), and *ɛ* is small constant set to 1*e −* 7 for numerical stability.

Following iBOT pretraining^31^ on ImageNet-22K, we pretrain a ViT-B/16 with 12 Transformer layers and a token size of 16*×*16 pixel. We use a batch size of 1,024 to process all training images, each with a resolution of 256*×*256 at 20*×* resized to 224*×*224 pixels. During training, the AdamW optimizer is used, and the learning rate is linearly ramped up over the initial five epochs to a base value of 5e-4, which is subsequently reduced by a cosine schedule to a final learning rate of 2e-6. Concurrently, the final layer of the network remains frozen during the first three epochs, while the teacher temperature incrementally ascends from 0.04 to 0.07 over a warm-up span of 30 epochs. Scaling of the two global crops is randomized within the range [0.32, 1.0], with the upper bound representing the utilization of the entire image, whereas values below this threshold indicate a proportional zoom, thereby employing only a segment of the original image. Masked Image Modeling (MIM) for these global crops is applied by selecting a masking ratio randomly sampled within the range of [0, 0.3], where a zero value signifies the complete absence of MIM. Values exceeding zero delineate the proportion of the global crops subject to masking. Additionally, a set of 10 local crops is rescaled accordingly by a randomly selected factor from the range [0.05, 0.32]. No masking is applied to the local crops. A detailed summary of hyperparameters used for pretraining is documented in **table S5**.

### Vision encoder fine-tuning

#### Patch annotations

Patch-level annotations were obtained through four distinct methodologies. First, publicly available annotations provided by Bayer Pharmaceuticals and Aignostics GmbH were employed^1^. They consist of polygonal annotations within 230 whole slide images from the TG-GATEs dataset. The annotations cover various lesions such as necrosis, mitosis, single-cell necrosis, extramedullary hematopoiesis, and cellular infiltration. These polygonal annotations were subsequently converted into patch-level annotations, where patches were retained if the original polygon overlapped with the patch coordinates. Manual inspection of each patch was applied to ensure label accuracy. In addition, semi-annotated human-in-the-loop annotations were generated using a weakly supervised slide classification system based on AttnPatchMIL. This method yielded pseudo patch labels that enabled the extraction of highly confident positive patches from the training set. A subsequent manual review led to the selection of true positive examples. To include normal patches, we extracted ten random patches from lesion-free slides, each thoroughly examined to exclude small lesions such as mitosis or single-cell necrosis that were not reported in the slide-level annotations. Lastly, human annotations were performed using the QuPath software^51^ to extract less common lesions, such as bile duct proliferation and fatty change. This process yielded a total of 29,795 patch annotations with lesions extracted from 3,539 different slides, and 13,888 normal patch annotations from 3,531 different slides (see **table S3**).

#### Model fine-tuning

TRACE was fine-tuned using all patch annotations from TG-GATEs train set. To do so, we employed a multilabel binary cross entropy objective with a class-stratified 80/20% train/validation split. The network was fine-tuned for 20 epochs with an initial learning rate of 4e-4 and layerwise learning decay of 0.65. Basic patch augmentations were performed during fine-tuning, based on random color jittering, mirroring, and rotation.

### Evaluation Setting

#### Vision encoder comparison

In our experiments, we compare TRACE against two pretrained vision encoders. First, a ResNet-50^37^ (8,543,296 parameters) pretrained on ImageNet^52^, where only the first three residual blocks are retained. This specific configuration is widely used in weakly supervised slide classification^53, 54^. We additionally compare TRACE against CTransPath^55^(28,289,038 parameters), a pretrained model based on a Swin Transformer^56^. CTransPath employs the “tiny” configuration with a window size of 14*×*14 pixels (Swin-T/14), and was pretrained on the TCGA and PAIP datasets^57^. CTransPath training was conducted using MoCoV3^58^for 100 epochs. In total, the pretraining process involves 15 million tissue patches and 32,120 WSIs. Both encoders use ImageNet image normalization parameters (mean and standard deviation).

#### Weakly-supervised slide classifiers

The weakly-supervised classification of slides is based on Multiple Instance Learning (MIL). MIL follows a two-stage pipeline. First, WSIs are tessellated into non-overlapping patches, which are further compressed using a pretrained patch embedding extractor, such as based on TRACE. Second, a learnable aggregator operator is trained to pool the patch embeddings into a single representation. A classifier further processes the slide-level representation to predict slide labels. In this work, all weakly supervised slide classification tasks are benchmarked against three MIL formulations: Attention-based MIL^35^, MeanMIL ^59^, and AttnPatchMIL (proposed).

#### Tissue segmentation and patching

Before MIL training, each slide was segmented with the CLAM toolbox^53^ that uses basic image processing tools to identify tissue regions from the background. After segmentation, non-overlapping 256 *×* 256-pixel patches were extracted at 20*×* magnification (0.49 *µ*m/px in TG-GATEs). Each patch was then resized to 224 *×* 224, normalized using default ImageNet mean and standard deviation parameters, and compressed using a pretrained vision encoder. To ensure a fair comparison, all the patch embeddings are extracted on the exact same set of patches.

#### Attention-based MIL

Attention-Based Multiple Instance Learning (ABMIL)^35^ is a widely employed MIL method in computational pathology. It uses an attention-based aggregator that assigns a weight to each patch embedding and then takes a weighted sum of the embeddings to derive a slide-level representation that can finally be used for classification. Specifically, we use a 2-layer Multilayer perceptron (MLP) pre-attention network with GeLU, LayerNorm, and dropout (0.1), followed by a gated attention mechanism with dropout (0.25) and Softmax normalization of the attention weights, followed by a 3-layer MLP classifier with GeLU activation and LayerNorm. All intermediate layers have 512 units, besides the 3-layer MLP classifier that uses 256 units.

#### MeanMIL

This baseline uses a simple aggregator that consists of the arithmetic mean of the patch embeddings ^59^. The resulting slide embedding is passed to a classifier (as in ABMIL).

#### AttnPatchMIL

This approach, which we primarily use for our study, uses a reformulation of ABMIL that enables jointly deriving patch and slide classification using slide-level supervision, at no additional cost. Specifically, each patch embedding is passed to a patch classification network (in this case, a multi-layer perceptron) and a gated attention network. Then, patch logits are weighted by the attention scores, which are finally summed to derive a slide-level prediction. As the slide-level prediction is directly connected to the patch-level logits, class-wise patch importance can be readily obtained without analyzing attention scores. AttnPatchMIL is implemented using 5-layer MLP patch classifier with LayerNorm, dropout (0.1) between all layers, and GELU activation. This formulation is similar to AdditiveMIL^60^, where the attention weights are scaling the patch logits instead of the patch embeddings.

All MIL models are trained using the RAdam optimizer^61^ with an initial learning rate of 1e-4, multi-label binary cross-entropy loss, and a maximum of 25 epochs with early stopping (patience set to 10) with respect to the validation loss (**table S6**).

#### Few-shot learning

We explore two approaches for learning with limited data and annotations: k-shot linear probing classification and one-shot visual prompting for morphological retrieval.

#### Linear probing classification

Following common practice in computer vision ^62–64^, we evaluate the performance of a logistic regression model (linear probing) in a low-data regime, where only a few examples per class are provided (few-shot learning). This setup enables assessing the representational power of the learned embeddings. Specifically, for a specific lesion, we sample *k* positive and *k* negative slides from TG-GATEs development set and obtain slide embeddings by averaging the patch embeddings within each slide. The examples are then used as a training dataset for the logistic regression module. All evaluations are performed on the entirety of TG-4k. We vary *k* from *k ∈ {*1, 5, 10, 25*}* to examine how the model benefits from more data points and repeat the experiments ten times to mitigate sampling biases.

#### Visual prompting

Visual prompting using one-shot learning is performed with two distinct prototyping approaches: single-patch prompting and single-slide prompting. In single-patch prompting, a single patch (256*×*256 pixel crop) displaying a specific lesion (such as basophilic cellular alteration) from a given study of interest is selected. Given the patch, the goal is to identify slides within the same study that contain the same morphology. On the other hand, the single-slide prompting uses an entire slide, instead of a patch, as a prompt. For a selected slide that contains a specific lesion of interest (positive slide), the slide prototype is defined by taking the arithmetic average of all patch embeddings within the slide. In both scenarios, the similarity between the prompt and a test slide is computed by first computing cosine similarity values between the prompt and test slide patch embeddings and then averaging the similarity within the test slide. Otsu thresholding method is applied to compute the threshold to determine whether the test slide is positive or negative.

#### Evaluation metrics

We report balanced accuracy, weighted F1 score, and macro-AUC for classification tasks. Studies are summarized with the log2 fold change. **Balanced accuracy** takes the class imbalance in the evaluation set into account by computing the unweighted average of the recall of each class. **Weighted F1 score** is computed by averaging the F1 score (the harmonic mean of precision and recall) of each class, further weighted by the respective support set size. **macro-AUC** is a threshold-free measure that computes the area under the receiver operating curve that plots the true positive rate against the false positive rate as the classification threshold changes. **Log2 fold change** is a measurement commonly used to quantify the relative change between two experimental conditions. It is calculated by taking the base 2 logarithm of the ratio between the percentage of a certain lesion under some conditions (such as high dose, sacrifice time of 29 days), and the percentage of that same lesion in the control group. To avoid artificially high Log2 fold change due to random noise in the predictions, we use a minimal threshold of 0.1% (i.e., all lesions that cover less than this size are ignored). Morphological Log2 fold change can be compared to fold change analyses widely employed in gene expression profiles to identify differentially expressed genes.

#### Statistical analysis

The reported error bars correspond to 95% confidence intervals derived using non-parametric bootstrapping using 100 bootstrap iterations. For all few-shot settings, we report results using box plots that indicate quartile values of model performance (*n* = 5 runs) with whiskers extending to data points within 1.5*×* the interquartile range.

### Reader study

The reader study comprised ten board-certified veterinary pathologists from three different countries (seven from Switzerland, two from the United States, and one from the United Kingdom), with an average of 10.2 years of experience post-board certification. Each pathologist participated in annotating an identical set of 100 slides randomly extracted from 22 studies.

#### Instructions to participants

Pathologists were asked to assign a severity score spanning normal, minimal, mild, moderate, and severe for every identified lesion. The pathologists underwent uniform training encompassing guidelines on employing a custom online slide viewer, along with 20 illustrative examples of regions of interest showcasing the target lesions. Each pathologist was directed to spend a maximum of 3 minutes per slide and to complete all 100 annotations within six weeks without collaborating with peers.

#### Defining consensus

After the reader study was conducted by all ten pathologists, three pathologists of the 10 pathologists with frequent exposure to rodent histopathology defined a consensus lesion scoring system and reviewed all 100 slides again based on this system. The final scores were then referred to as the gold standard and used for comparing each pathologist and the AI system. First, all lesions were rigorously defined and characterized as per the INHAND guidelines (International Harmonization of Nomenclature and Diagnostic Criteria, see **table S2**). Next, lesion scores were defined based on the proposed INHAND scoring system as, where minimal is INHAND score 1, mild is 2, moderate is 3, and severe is 4 and 5. For singlecell necrosis and increased mitosis, a tailored scoring was defined where individual single-cell necrosis, respectively mitosis, were manually counted in ten (400x) high power fields, starting in the hot spot area.

#### From quantification scores to severity scores

TRACE (FT) outputs quantification scores that describe the size of the tissue with lesions normalized by the total tissue size. For instance, a quantification score of 2.0 means that 2.0% of the tissue is necrotic. To compare quantification scores of TRACE (FT) with the consensus (i.e., a score from normal to minimal to mild to moderate to severe), we need to convert them into severity scores. To this end, we employ leave-one-out cross-validation, where we iteratively predict the severity of each lesion in each slide from the consensus scores. In this way, we learn a mapping between consensus and quantification scores without any form of data leakage. This further allows to compute Quadratic Cohen’s score to rigorously compare the consensus with TRACE (FT).

### Computing hardware and software

All experiments were conducted using Python (version 3.9), deep learning components were implemented using PyTorch (version 2.0, CUDA version 11.7). We used the original implementation of the iBOT model (github.com/bytedance/ibot) for training the vision encoder. WSI pre-processing and manipulation was done using OpenSlide (version 4.3.1) and openslide-python (version 1.2.0). We used Scikit-learn (version 1.2.1) and TorchMetrics (version 1.1.0) to evaluate the performance of the proposed methods and baselines. ResNet50 weights pretrained on ImageNet were downloaded from TorchVision (version 0.15), CTransPath weights and implementation was adapted from the original implementation (https://github.com/Xiyue-Wang/TransPath). Pandas (1.4.2), Numpy (1.21.5), Pillow (version 9.3.0) and OpenCV-python (3.3.1) were used to perform basic image and array processing tasks. Matplotlib (version 3.7) was used to generate plots and figures. Low-dimensional representations of TRACE (FT) were obtained using the python package Uniform Manifold Approximation and Projection (UMAP) version 0.5.6. The iBot pretraining was done on 8*×*80GB NVIDIA A100 GPUs configured for multi-GPU training using distributed data-parallel (DDP) (pytorch.org). Downstream experiments were conducted on 3*×* 24GB NVIDIA 3090 GPUs. Slides were annotated and visualized using QuPath (version 0.4.3). The viewer used for the reader study and the online demo is based on OpenSeadragon (version 4.1.0) and JavaScript (version ES13).

## Data availability

TG-GATEs data consisting of whole-slide images and slide labels can be freely accessed from the National Institute of Biomedical Innovation portal ^2^. Pixel annotations on a subset of 230 TG-GATEs can be accessed freely from Zenodo (https://zenodo.org/record/7541930) Patch annotations can be shared on a case-by-case basis based on needs. Pseudo-patch annotations obtained by running the fine-tuned patch encoder can be shared on a case-by-case basis based on needs.

## Code availability

Upon publication, the authors will release code for extracting features using TRACE and for performing the weakly-supervised evaluation. In addition, authors will release the pretrained TRACE patch encoder along with its fine-tuned version.

## Author contributions

G.J. and S.B. conceived the study and designed the experiments. G.J., A.S., D.W., L.O., A.Z., R.J.C., R.P, and L.L. ran experiments and provided experimental feedback. S.B., J.A., S.B., M.D., C.G, L.R., S.S., S.K., S.R., J.D. participated in the reader study. G.J. prepared the manuscript with input from all co-authors. F.M. supervised the research.

## Acknowledgements

This work was supported in part by the BWH & MGH Pathology, BWH president’s fund, Massachusetts Life Sciences Center, NIGMS R35GM138216 (F.M.), and the BWH President’s Scholar fund (G.G.). R.J.C. was also supported by the NSF Graduate Fellowship. L.O. was supported by the German Academic Exchange (DAAD) Fellowship. We thank Timothy Janicki, Richard Kenny, and the system administration staff at the MGB Enterprise Research Infrastructure & Services (ERIS) Research Computing Core for their dedicated support in providing and maintaining access to computing resources. We warmly thank Dr. Pierre Moulin for the early-stage discussions on toxicity assessment and computational toxicology.

---

**Figure S1:**
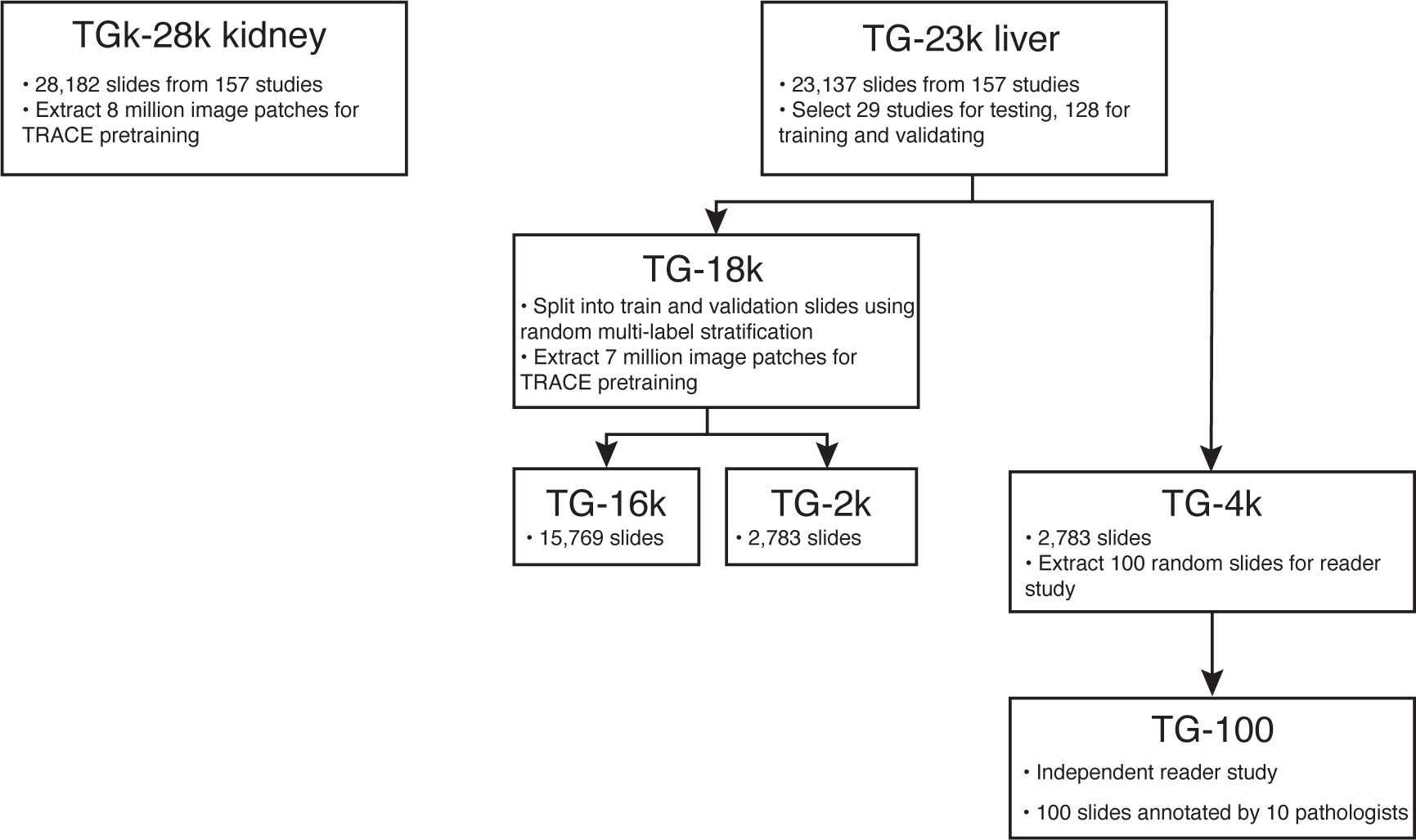
Overview of data profile. This study uses the TG-GATEs database for TRACE pretraining (TG-28k, TG-18k), weakly supervised and few-shot classification (TG-4k), TRACE fine-tuning with patch annotations, and conducting the reader study (TG-100).

**Figure S2:**
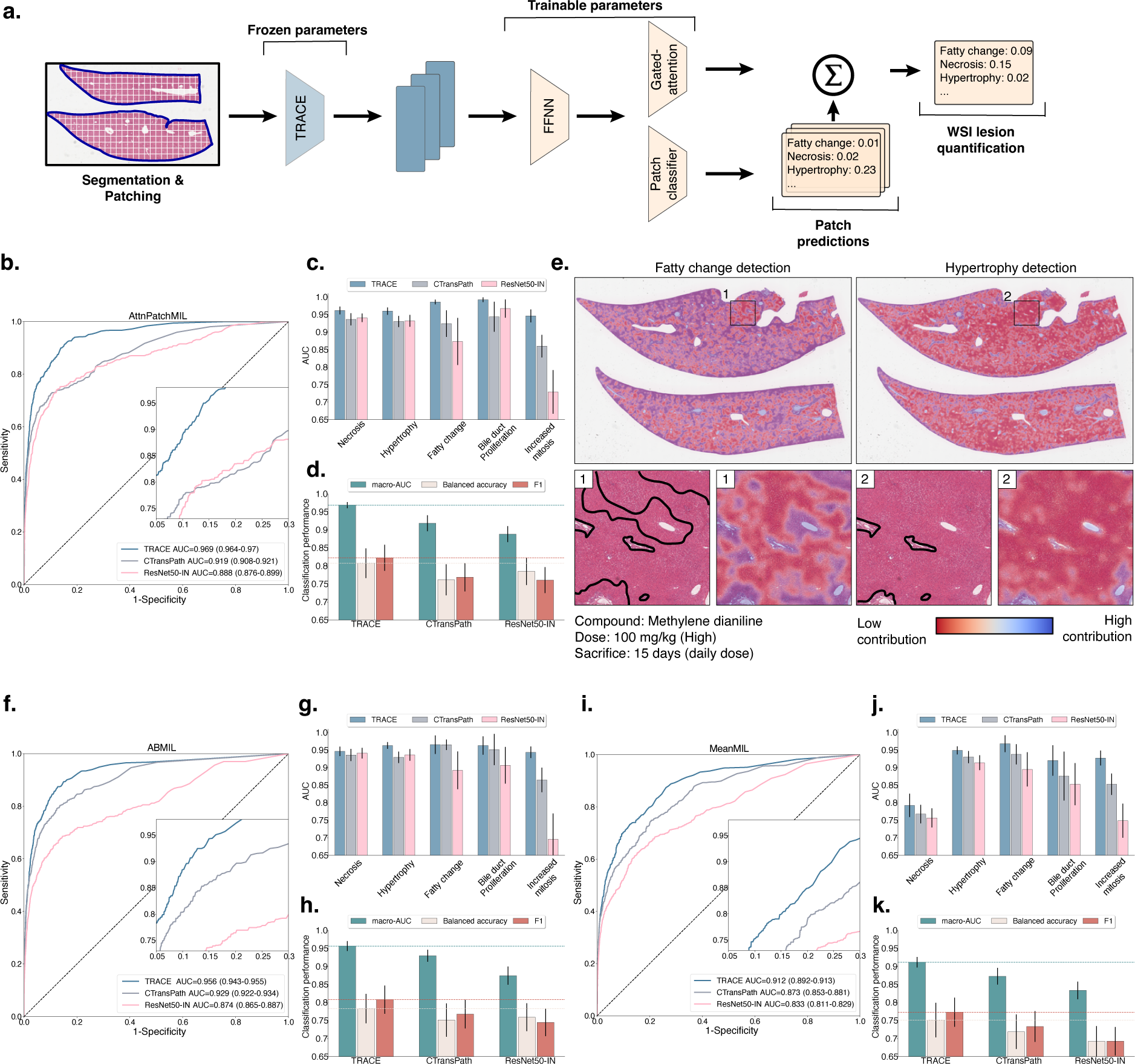
Weakly supervised slide classification. **a.** Overview of the proposed multiple instance learning (MIL) architecture, AttnPatchMIL, for joint patch and slide classification using slide-level supervision. **b,c,d.** Evaluation of AttnPatchMIL on 5-class lesion classification comparing TRACE against ResNet50-IN and CTransPath vision encoders. Evaluation based on multi-label ROC and macro-AUC in **b.**; class-wise AUC in **c.** and; overall balanced accuracy, macro-AUC, and F1 in **d.**. **e.** Patch-wise attribution of fatty change and hypertrophy using AttnPatchMIL. **f.,g.,h.** Analogous evaluation using Attention-based MIL (ABMIL). **i.,j.,k.** Analogous evaluation using MeanMIL based on the mean patch embedding. All models are trained on TG-18k and tested on TG-4k. Error bars in **b,c,d,f,g,h,i,j,k.** represent 95% confidence intervals and were computed using non-parametric bootstrapping (100 iterations). ROC: receiver operating characteristic; AUC: area under the ROC curve; FFNN: feed-forward neural network.

**Figure S3:**
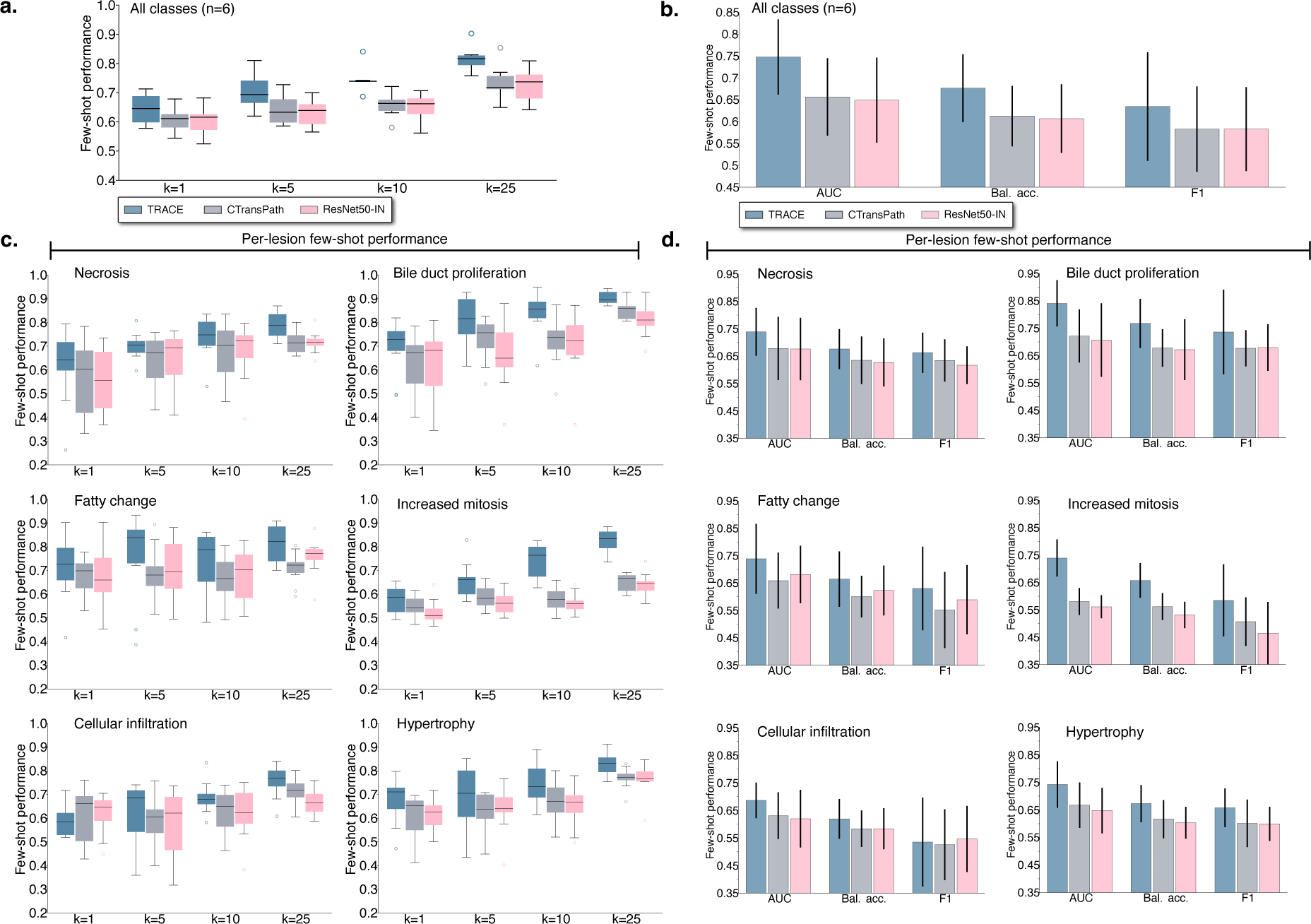
Few-shot classification. **a.** Comparison of TRACE, CTransPath and ResNet50-IN vision encoders for few-shot learning (*k ∈* 1, 5, 10, 25) in TG-4k evaluated AUC averaged across six binary classification tasks. **b.** Comparison of TRACE, CTransPath and ResNet50-IN vision encoders for *k* =10 in TG-4k evaluated using macro-AUC, balanced accuracy and F1 score. **c.** Per-class (n=6 lesions) few-shot performance evaluated using macro AUC. **d.** Per-class (n=6 lesions) few-shot performance evaluated using macro AUC with *k* =10. Error bars in **a,c.** represent 95% confidence intervals and were computed using non-parametric bootstrapping (100 iterations). Error bars in **b,d.** represent standard deviation and were computed using classification performance repeated over 10 runs, where each run samples a different random set of *k* training samples per class. AUC: area under the ROC curve; Bal. acc.: balanced accuracy.

**Figure S4:**
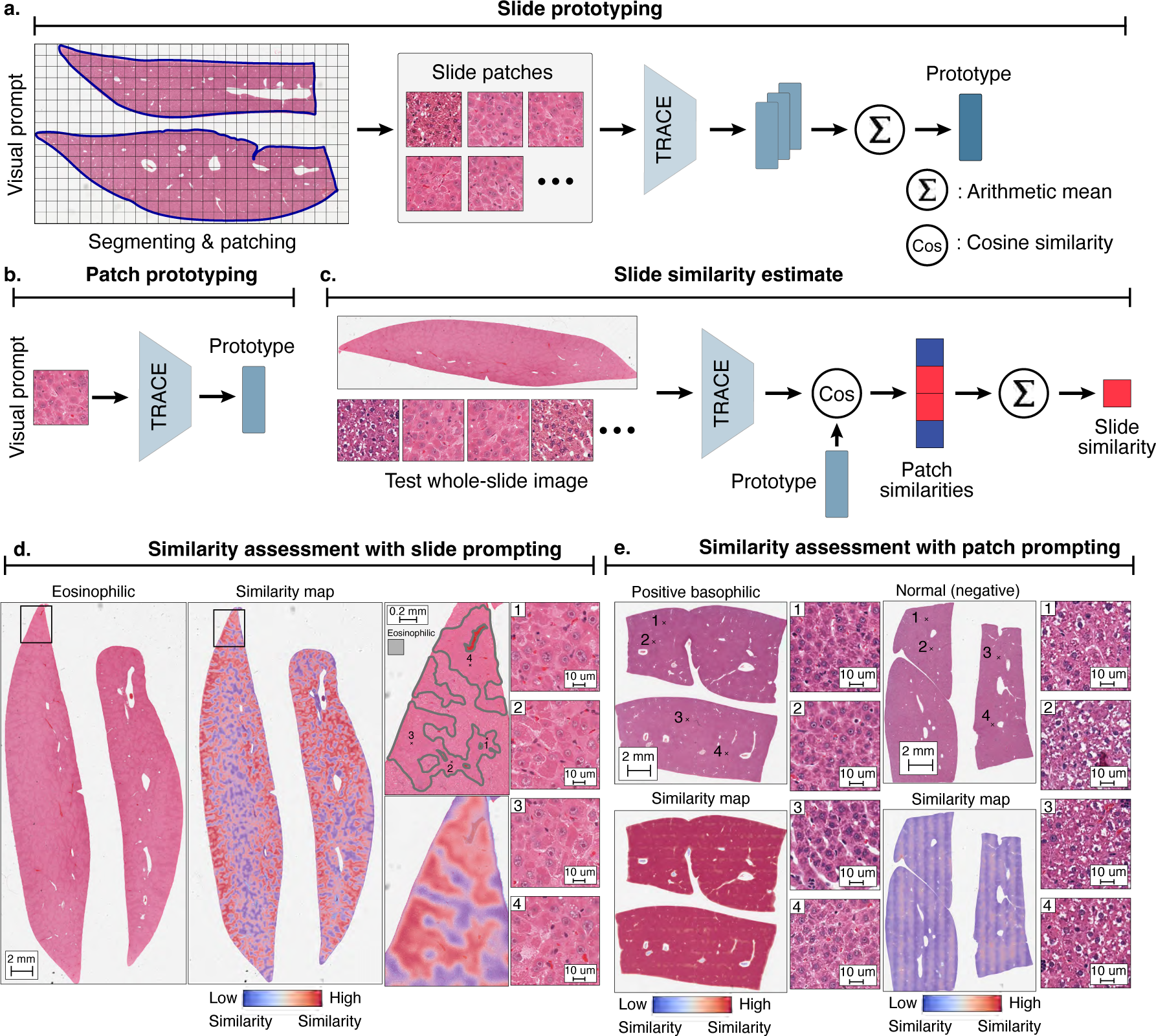
Patch and slide-level prototyping. **a.** A slide-level prototype is defined by taking the average TRACE patch embeddings of a slide that contains a distinct morphology of interest. **b.** A patch-level prototype is defined as a single TRACE patch embedding that contains morphology of interest. **c.** A similarity score is defined by computing the average cosine distance between the prototype (patch or slide) and all patch embeddings from a test slide, with the binary prediction (positive if the slide contains morphology of interest, and negative otherwise) threshold determined by Otsu method. **d.** Similarity assessment using single-slide prompting for detecting eosinophilic cellular alteration in thioacetamide and basophilic cellular alteration in puromycin aminonucleoside. Visualization of the patch-level similarity with the prototype yields a pseudo-segmentation map indicating the presence of the morphology of interest. In a high-dose thioacetamide slide, annotations of eosinophilic regions align almost perfectly with the patch-level similarity. **e.** Similarity assessment using single-patch prompting classification of basophilic cellular alteration. High similarity is identified between the positive basophilic prompt and the positive test slide (center), and low similarity with the negative (normal) slide (right).

**Figure S5:**
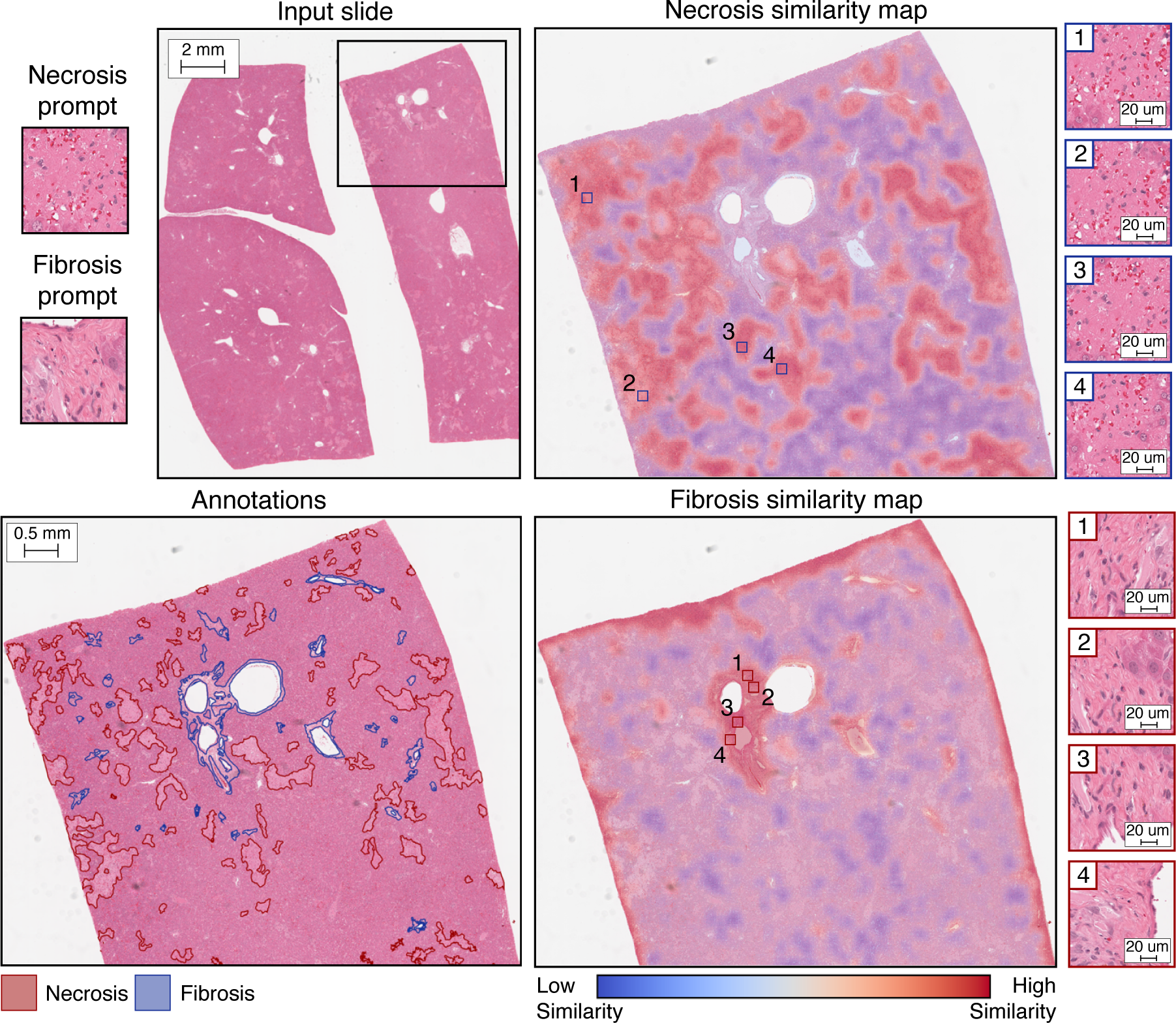
Similarity assessment with single-patch prompting. Detection of necrosis and fibrosis using single-patch prompting upon daily administration (15 days) of 30 mg/kg of monocrotaline.

**Figure S6:**
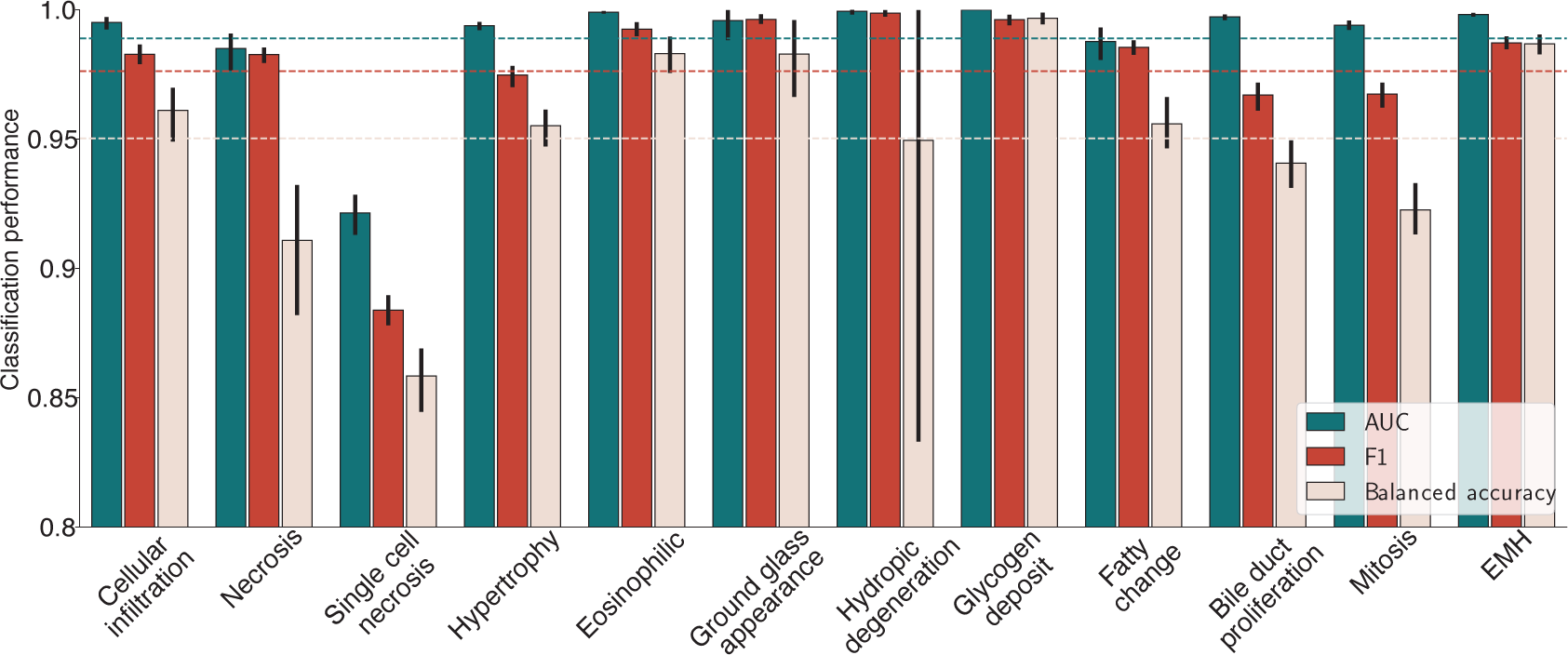
Patch classification evaluation. Class-wise macro-AUC, balanced accuracy and F1 score of TRACE (FT). Error bars represent 95% confidence intervals and were computed using non-parametric bootstrapping (100 iterations). EMH: extramedullary hematopoiesis; Eosinophilic: eosinophilic cellular alteration.

**Figure S7:**
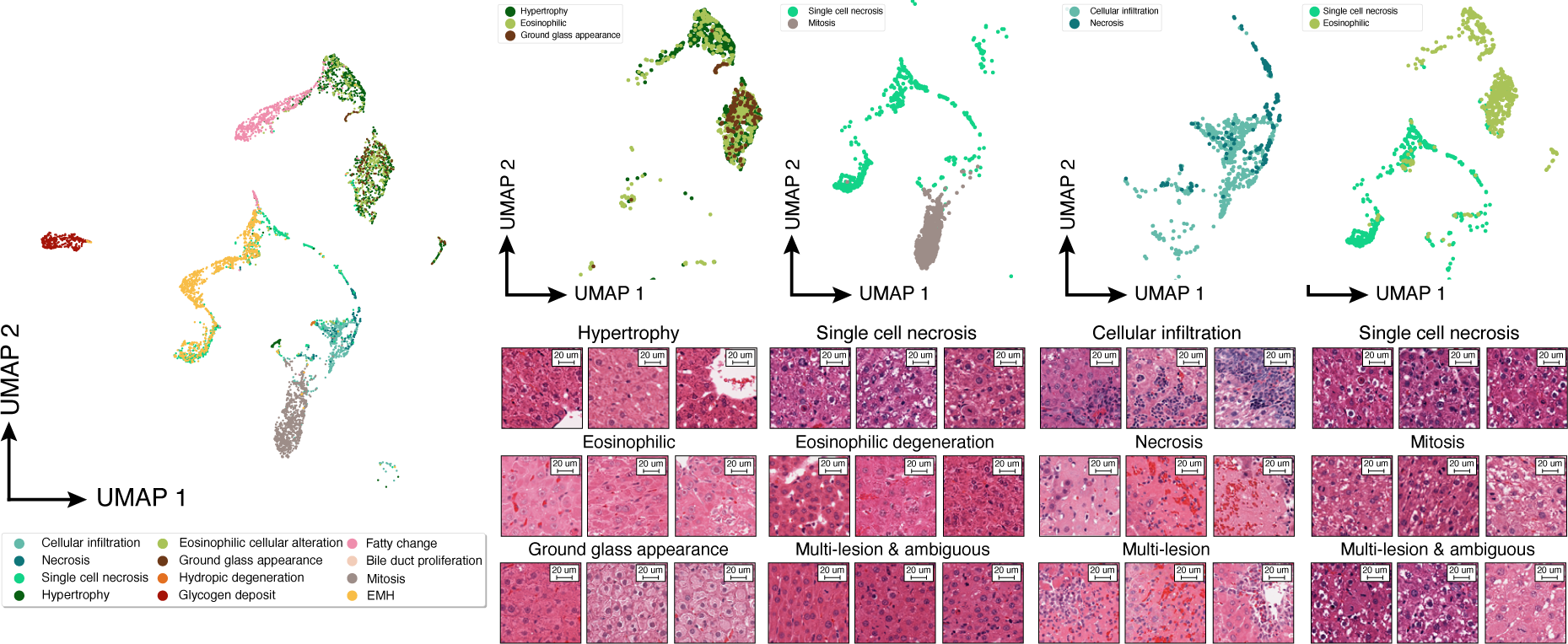
Visualization of TRACE (FT) embedding space. Uniform Manifold Approximation and Projection (UMAP) of TRACE (FT) patch embedded colored by their annotation. All shown patches are from TG-4k. Zoom on specific regions of the latent space with patch examples. Each patch is 256*×*256 pixels at 20*×*. EMH: extramedullary hematopoiesis. Eosinophilic: eosinophilic cellular alteration.

**Figure S8:**
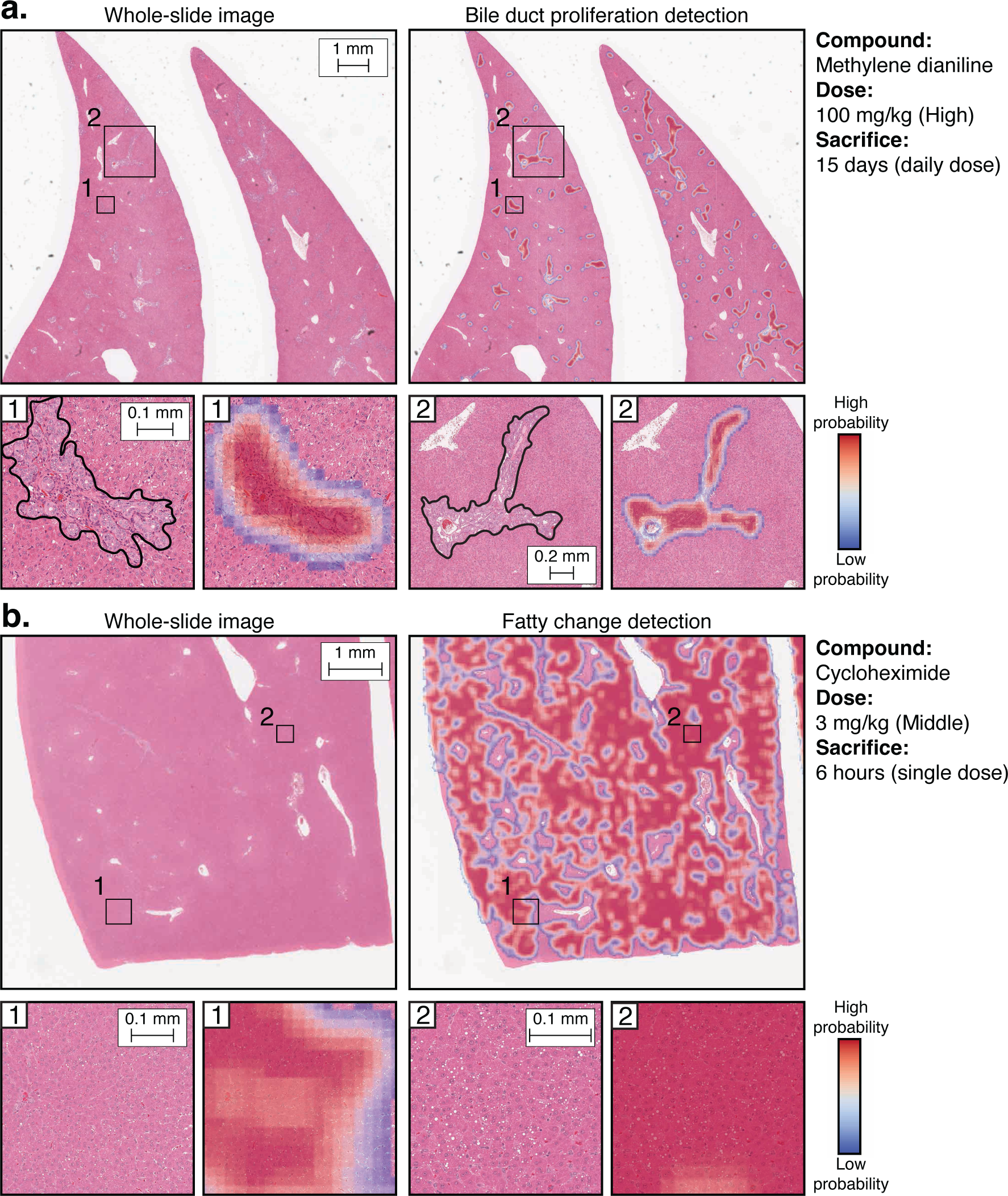
Visualization of TRACE fine-tuned with patch annotations. **a.** Example of lesion detection and segmentation using TRACE (FT) in a high dose sample administered with methylene dianiline. Regions highlight bile duct proliferation. **b.** Example of lesion detection and segmentation using TRACE (FT) in a middle dose sample administered with cycloheximide. Regions highlight hepatocellular fatty change. All results were obtained using 80% patch overlap.

**Figure S9:**
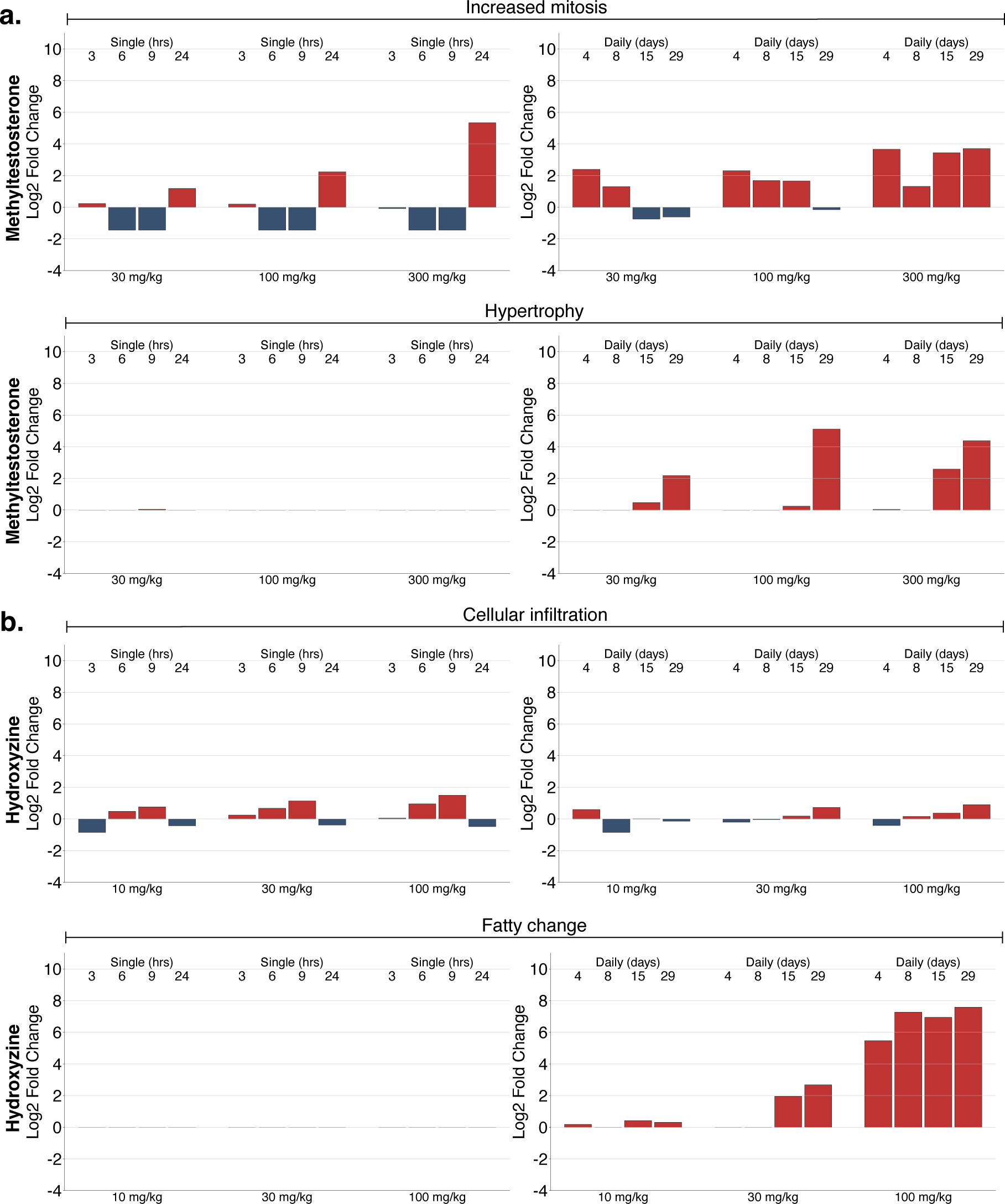
Automatic dose-response toxicity assessment. **a.** Morphological lesion log2 fold change of increased mitosis and hypertrophy in methyltestosterone in single and daily dose sample groups with respect to the control group. **b.** Log2 fold change of cellular infiltration and fatty change in hydroxyzine in single and daily doses with respect to the control group. Plots without bars indicate that no lesion was detected.

**Table S1:** Distribution of lesions used in weakly supervised slide classification. TG-GATEs includes 23,136 liver WSIs from 157 different pre-clinical studies. Necrosis refers to single-cell, focal/multifocal, or zonal hepatocellular necrosis; hypertrophy refers to enlarged hepatocytes, primarily due to an increase in the cytosolic protein or number of organelles; fatty change includes macro and microvesicular hepatocellular vacuolation; bile duct proliferation refers to bile duct hyperplasia with an increased number of bile ducts; we additionally include oval cell proliferation. Increased mitosis refers to hepatocyte mitosis above normal levels. Complementary information describing each lesion is provided in **table S2** WSIs, whole-slide images.

| Lesions | Train/Validation |  | Test |  |
| --- | --- | --- | --- | --- |
|  | WSIs | Compounds | WSIs | Compounds |
| Necrosis | 748 | 101 | 264 | 17 |
| Hypertrophy | 1649 | 109 | 331 | 18 |
| Fatty change | 308 | 33 | 73 | 7 |
| Bile duct proliferation | 111 | 10 | 41 | 2 |
| Increased mitosis | 331 | 43 | 94 | 9 |

**Table S2:** Lesion definition. Definitions and scope are based on the INHAND guidelines^40^. INHAND is the International Harmonization of Nomenclature and Diagnostic Criteria, a publicly accessible resource that defines guidelines to diagnose lesions found in rodent toxicity and carcinogenicity studies. When INHAND lacked specific guidelines regarding diagnosing, such as for scoring “increased mitosis”, we relied on the National Toxicology Program guidelines available online https://ntp.niehs.nih.gov/atlas/nnl/hepatobiliary-system/liver. WSC: weakly supervised classification; FSL: few-shot learning; MR: morphological retrieval; PC: patch classification, RS: reader study.

| Lesion | Definition | Used in |
| --- | --- | --- |
| *Hepatocellular responses, cellular degeneration, injury, and death* |  |  |
| Fatty change | Hepatocellular vacuolation, consistent with intracytoplasmic lipid accumulation. Includes macro and microvesicular. | WSC, FSL, PC, RS |
| Hydropic degeneration | Hepatocellular vacuolation, consistent with intracytoplasmic fluid accumulation. | PC |
| Necrosis | Cell death of hepatocytes. Includes focal/multifocal and zonal (centrilobular, midzonal, periportal and diffuse). | WSC, FSL, MR, PC, RS |
| Single-cell necrosis | Necrosis (or apoptosis) of single hepatocytes. | PC, RS |
| Hypertrophy | Enlargement of the hepatocyte cytoplasm, secondary to increase in the cytosolic protein or number of organelles. | WSC, FSL, PC, RS |
| Glycogen deposit | Hepatocellular cytoplasmic alteration, consistent with glycogen accumulation. | PC, RS |
| *Inflammatory cell infiltrates, inflammatory cell infiltrates and hepatic* |  |  |
| Cellular infiltration | Infiltrations of inflammatory cells in the liver. Includes neutrophil, mononuclear and peribiliary. | FSL, PC, RS |
| Fibrosis | The presence of fibrous connective tissue in the liver above normal levels. Includes pericellular, peribiliary, and postnecrotic fibrosis. | MR |
| *Non-neoplastic proliferative lesions* |  |  |
| Basophilic cellular alteration | Hepatocellular cytoplasmic basophilia, due to free ribosomes or rough endoplasmic reticulum. | MR |
| Eosinophilic cellular alteration | Hepatocellular cytoplasmic eosinophilia, due to an increase in cytoplasmic organelles. | PC |
| Ground glass appearance | Hepatocytes with glassy and hypereosinophilic appearing cytoplasm | PC |
| Bile duct proliferation | Increased number of small bile ducts arising in portal region. Biliary epithelium appears normal or may show degenerative or atrophic changes. | WSC, FSL, PC, RS |
| Increased mitosis | Increased hepatocyte mitoses above normal background levels (>1-2 mitotic figures per 10 (400x) HPF). | WSC, FSL, PC, RS |
| *Other lesions* |  |  |
| Extramedullary hematopoiesis | Hematopoietic cell proliferation in the liver. Aggregates of hematopoietic cells are distributed in the hepatic sinusoids as well as around central veins and portal areas. | PC, RS |

**Table S3:**
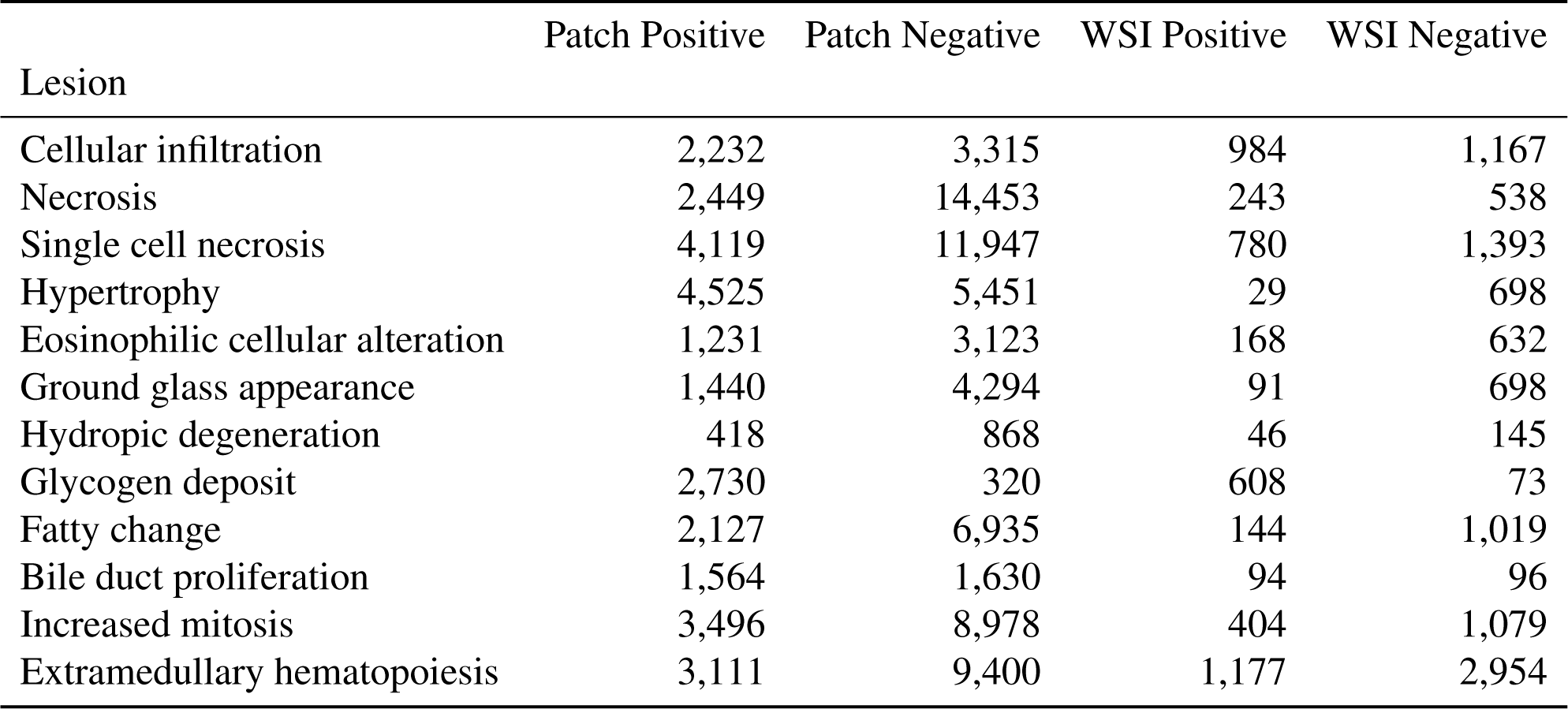
Distribution of patch annotations. As many patches include several labels, such as fatty change and hypertrophy, we report positive patches (lesion is present) and the number of hard negatives (lesion is not present). Positive patches may include more than one lesion.

| Lesion | Patch Positive | Patch Negative | WSI Positive | WSI Negative |
| --- | --- | --- | --- | --- |
| Cellular infiltration | 2,232 | 3,315 | 984 | 1,167 |
| Necrosis | 2,449 | 14,453 | 243 | 538 |
| Single cell necrosis | 4,119 | 11,947 | 780 | 1,393 |
| Hypertrophy | 4,525 | 5,451 | 29 | 698 |
| Eosinophilic cellular alteration | 1,231 | 3,123 | 168 | 632 |
| Ground glass appearance | 1,440 | 4,294 | 91 | 698 |
| Hydropic degeneration | 418 | 868 | 46 | 145 |
| Glycogen deposit | 2,730 | 320 | 608 | 73 |
| Fatty change | 2,127 | 6,935 | 144 | 1,019 |
| Bile duct proliferation | 1,564 | 1,630 | 94 | 96 |
| Increased mitosis | 3,496 | 8,978 | 404 | 1,079 |
| Extramedullary hematopoiesis | 3,111 | 9,400 | 1,177 | 2,954 |

**Table S4:** Cases used in the reader study. A total of 100 tissue sections were randomly extracted from TG-GATEs test set (TG-4k).

| Slide ID | Compound | Dose | Single or Repeat | Sacrifice |
| --- | --- | --- | --- | --- |
| 10921 | Griseofulvin | 1000 | Single | 3 hrs |
| 11089 | Griseofulvin | 1000 | Single | 24 hrs |
| 13326 | Metformin | 0 | Single | 9 hrs |
| 14012 | Methyldopa | 200 | Single | 3 hrs |
| 14430 | Methyldopa | 0 | Repeat | 15 days |
| 14478 | Methyldopa | 60 | Repeat | 15 days |
| 19714 | Hydroxyzine | 30 | Repeat | 15 days |
| 20367 | Mefenamic acid | 300 | Repeat | 8 days |
| 20422 | Mefenamic acid | 300 | Repeat | 15 days |
| 27539 | Thioacetamide | 0 | Single | 6 hrs |
| 30270 | Bromobenzene | 300 | Single | 24 hrs |
| 30272 | Bromobenzene | 300 | Single | 24 hrs |
| 30274 | Bromobenzene | 300 | Single | 24 hrs |
| 31778 | Cyclophosphamide | 15 | Repeat | 15 days |
| 33012 | Metformin | 300 | Repeat | 4 days |
| 33056 | Metformin | 1000 | Repeat | 4 days |
| 33057 | Metformin | 1000 | Repeat | 4 days |
| 46078 | Ethionamide | 300 | Single | 9 hrs |
| 46360 | Ethionamide | 100 | Repeat | 4 days |
| 48472 | Phenacetin | 0 | Single | 24 hrs |
| 48744 | Phenacetin | 2000 | Single | 6 hrs |
| 48936 | Phenacetin | 100 | Repeat | 4 days |
| 49066 | Phenacetin | 1000 | Repeat | 8 days |
| 50226 | Danazol | 100 | Repeat | 15 days |
| 50987 | Cisplatin | 0 | Single | 6 hrs |
| 51434 | Cisplatin | 1 | Repeat | 4 days |
| 51604 | Carboplatin | 0 | Single | 6 hrs |
| 53169 | Bromoethylamine | 6 | Repeat | 29 days |
| 53257 | Bromoethylamine | 20 | Repeat | 29 days |
| 53335 | Mexiletine | 40 | Single | 9 hrs |
| 54187 | Cephalothin | 1000 | Single | 3 hrs |
| 54409 | Cyclosporine a | 0 | Repeat | 4 days |
| 54597 | Cyclosporine a | 10 | Repeat | 15 days |
| 54816 | Cyclosporine a | 100 | Repeat | 8 days |
| 54822 | Cyclosporine a | 100 | Repeat | 15 days |

Table S4: Continuation of table S4.
| Slide ID | Compound | Dose | Single or Repeat | Sacrifice |
| --- | --- | --- | --- | --- |
| 55649 | Gentamicin | 30 | Repeat | 8 days |
| 57174 | Danazol | 1000 | Single | 24 hrs |
| 57236 | Danazol | 2000 | Single | 6 hrs |
| 57336 | Theophylline | 0 | Single | 6 hrs |
| 58909 | Cycloheximide | 10 | Single | 3 hrs |
| 59123 | Tunicamycin | 300 | Single | 3 hrs |
| 6009 | Diazepam | 0 | Single | 3 hrs |
| 6013 | Diazepam | 0 | Single | 6 hrs |
| 60435 | Isoniazid | 2000 | Single | 6 hrs |
| 62014 | Hexachlorobenzene | 0 | Single | 3 hrs |
| 6224 | Hexachlorobenzene | 0 | Repeat | 15 days |
| 63748 | Methylene dianiline | 30 | Repeat | 8 days |
| 63941 | Methylene dianiline | 100 | Repeat | 29 days |
| 63952 | Methylene dianiline | 100 | Repeat | 29 days |
| 63955 | Methylene dianiline | 100 | Repeat | 29 days |
| 10913 | Griseofulvin | 300 | Single | 3 hrs |
| 10969 | Griseofulvin | 300 | Single | 6 hrs |
| 11080 | Griseofulvin | 1000 | Single | 24 hrs |
| 14008 | Methyldopa | 200 | Single | 3 hrs |
| 14023 | Methyldopa | 200 | Single | 6 hrs |
| 14211 | Methyltestosterone | 300 | Single | 24 hrs |
| 14554 | Methyldopa | 200 | Repeat | 29 days |
| 19318 | Hydroxyzine | 10 | Single | 24 hrs |
| 19764 | Hydroxyzine | 100 | Repeat | 15 days |
| 19773 | Hydroxyzine | 100 | Repeat | 15 days |
| 19777 | Hydroxyzine | 100 | Repeat | 29 days |
| 19781 | Hydroxyzine | 100 | Repeat | 29 days |
| 19785 | Hydroxyzine | 100 | Repeat | 29 days |
| 20370 | Mefenamic acid | 300 | Repeat | 8 days |
| 20414 | Mefenamic acid | 300 | Repeat | 15 days |
| 20477 | Mefenamic acid | 300 | Repeat | 29 days |
| 20483 | Mefenamic acid | 300 | Repeat | 29 days |
| 2421 | Isoniazid | 100 | Single | 6 hrs |
| 2589 | Isoniazid | 0 | Repeat | 4 days |
| 30278 | Bromobenzene | 300 | Single | 24 hrs |

Table S4: Continuation of table S4.
| Slide ID | Compound | Dose | Single or Repeat | Sacrifice |
| --- | --- | --- | --- | --- |
| 30465 | Bromobenzene | 30 | Repeat | 15 days |
| 33006 | Metformin | 100 | Repeat | 29 days |
| 33077 | Metformin | 1000 | Repeat | 15 days |
| 46792 | Thioacetamide | 45 | Repeat | 4 days |
| 48569 | Phenacetin | 300 | Single | 9 hrs |
| 48685 | Phenacetin | 1000 | Single | 9 hrs |
| 48998 | Phenacetin | 100 | Repeat | 15 days |
| 49064 | Phenacetin | 1000 | Repeat | 8 days |
| 51220 | Cisplatin | 3 | Single | 24 hrs |
| 52036 | Carboplatin | 0 | Repeat | 15 days |
| 53036 | Bromoethylamine | 2 | Repeat | 29 days |
| 53941 | Cephalothin | 0 | Single | 3 hrs |
| 53997 | Cephalothin | 0 | Single | 9 hrs |
| 54300 | Cephalothin | 2000 | Single | 3 hrs |
| 54937 | Puromycin aminonucleoside | 0 | Single | 9 hrs |
| 54991 | Gentamicin | 0 | Single | 24 hrs |
| 55355 | Puromycin aminonucleoside | 120 | Single | 6 hrs |
| 55696 | Gentamicin | 30 | Repeat | 15 days |
| 55728 | Puromycin aminonucleoside | 40 | Repeat | 8 days |
| 56895 | Danazol | 0 | Single | 6 hrs |
| 57802 | Theophylline | 0 | Repeat | 8 days |
| 58319 | Acetazolamide | 200 | Single | 9 hrs |
| 58819 | Cycloheximide | 3 | Single | 6 hrs |
| 59224 | Tunicamycin | 300 | Single | 24 hrs |
| 60413 | Isoniazid | 200 | Single | 6 hrs |
| 60648 | Cyclophosphamide | 50 | Single | 9 hrs |
| 62091 | Hexachlorobenzene | 1000 | Single | 6 hrs |
| 6302 | Hexachlorobenzene | 30 | Repeat | 29 days |
| 63875 | Methylene dianiline | 100 | Repeat | 15 days |
| 70760 | Phenacetin | 1000 | Repeat | 8 days |

**Table S5:** iBOT hyperparameters used in TRACE pretraining. The training converged after 80 epochs for a total training time of 208 hours using 8 *×* 80GB NVIDIA A100 GPUs.

| Hyper-parameter | Value |
| --- | --- |
| Layers | 12 |
| Heads | 12 |
| Patch size | 16 |
| FFN layer | MLP |
| Head activation | GELU |
| Embedding dimension | 768 |
| Stochastic dropout rate | 0.1 |
| Global crop scale | 0.32, 1.0 |
| Global crop number | 2 |
| Local crop scale | 0.05, 0.32 |
| Local crop number | 10 |
| Max masking ratio | 0.3 |
| Min masking ratio | 0.0 |
| Gradient clipping max norm | 0.3 |
| Normalize last layer | ✓ |
| Shared head | ✓ |
| head output dimension | 8192 |
| Optimizer | AdamW |
| Batch size | 1024 |
| Freeze last layer (it, ep) | 44124, 3 |
| Warmup (it, ep) | 73540, 5 |
| Warmup teacher temperature (it, ep) | 441240, 30 |
| Max training (it, ep) | 1176640, 80 |
| Number of images | 15061790 |
| Learning rate schedule | Cosine |
| Learning rate (start) | 0.0 |
| Learning rate (post warmup) | 0.0005 |
| Learning rate (final) | 0.000002 |
| Teacher temperature (start) | 0.04 |
| Teacher temperature (final) | 0.07 |
| Teacher momentum | 0.996 |
| Weight decay (start) | 0.04 |
| Weight decay (end) | 0.4 |
| Automatic mixed precision | fp16 |

**Table S6:**
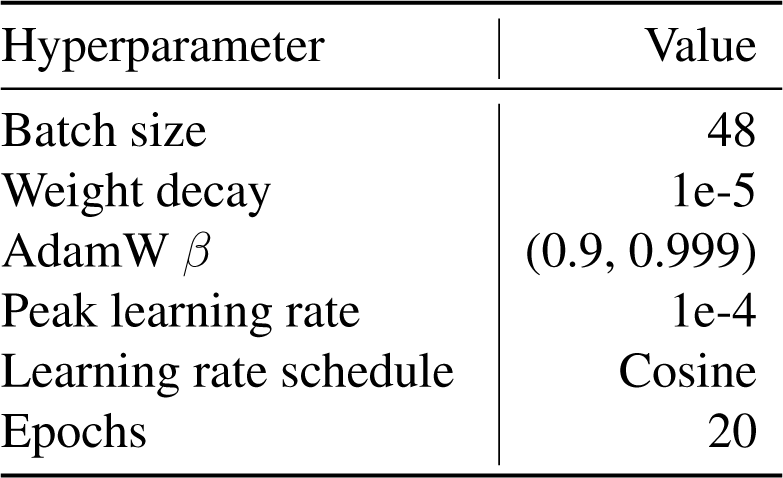
Hyperparameters used in slide-level supervised classification. A single 24GB NVIDIA GeForce RTX 3090 GPU was used to train the multiple instance learning models.

| Hyperparameter | Value |
| --- | --- |
| Batch size | 48 |
| Weight decay | 1e-5 |
| AdamW $\beta$ | (0.9, 0.999) |
| Peak learning rate | 1e-4 |
| Learning rate schedule | Cosine |
| Epochs | 20 |

## Footnotes

1 https://dbarchive.biosciencedbc.jp/en/open-tggates/download.html

1 https://zenodo.org/record/7541930

2 https://dbarchive.biosciencedbc.jp/en/open-tggates/desc.html

## References

1. Seyhan, A. A. Lost in translation: the valley of death across preclinical and clinical divide – identification of problems and overcoming obstacles. Translational Medicine Communications 4, 1–19 (2019).

2. Cook, D. et al. Lessons learned from the fate of AstraZeneca’s drug pipeline: a five-dimensional framework. Nat. Rev. Drug Discovery 13, 419–431 (2014).

3. Waring, M. J. et al. An analysis of the attrition of drug candidates from four major pharmaceutical companies. Nat. Rev. Drug Discovery 14, 475–486 (2015).

4. U.S. Food and Drug Administration. 21 CFR Part 58 – Good Laboratory Practice for Nonclinical Laboratory Studies (2024). URL https://www.ecfr.gov/current/title-21/chapter-I/subchapter-A/part-58. [Online; accessed 4. Mar. 2024].

5. Patel, M., Taskar, K. S. & Zamek-Gliszczynski, M. J. Importance of Hepatic Transporters in Clinical Disposition of Drugs and Their Metabolites. J. Clin. Pharmacol. 56, S23–S39 (2016).

6. Weaver, R. J. et al. Managing the challenge of drug-induced liver injury: a roadmap for the development and deployment of preclinical predictive models. Nat. Rev. Drug Discovery 19, 131–148 (2020).

7. Chen, M. et al. Dilirank: the largest reference drug list ranked by the risk for developing drug-induced liver injury in humans. Drug Discovery Today 21, 648–653 (2016).

8. Morton, D. et al. Recommendations for pathology peer review. Toxicologic Pathology 38, 1118–1127 (2014).

9. Schafer, K. A. et al. Use of severity grades to characterize histopathologic changes. Toxicologic Pathology 46, 256–265 (2018).

10. Serag, A. et al. Translational ai and deep learning in diagnostic pathology. Frontiers in Medicine 6, 185 (2019).

11. LeCun, Y., Bengio, Y. & Hinton, G. Deep learning. Nature 521, 436–444 (2015).

12. Madabhushi, A. & Lee, G. Image analysis and machine learning in digital pathology: Challenges and opportunities. Medical image analysis 33, 170–175 (2016).

13. Van der Laak, J., Litjens, G. & Ciompi, F. Deep learning in histopathology: the path to the clinic. Nature Medicine 27, 775–784 (2021).

14. Shmatko, A., Ghaffari Laleh, N., Gerstung, M. & Kather, J. N. Artificial intelligence in histopathology: enhancing cancer research and clinical oncology. Nature Cancer 3, 1026– 1038 (2022).

15. Song, A. H. et al. Artificial intelligence for digital and computational pathology. Nature Reviews Bioengineering (2023).

16. Turner, O. C. et al. Society of toxicologic pathology digital pathology and image analysis special interest group article*: Opinion on the application of artificial intelligence and machine learning to digital toxicologic pathology. Toxicologic Pathology 48, 277–294 (2020).

17. Hoefling, H. et al. Histonet: A deep learning-based model of normal histology. Toxicologic Pathology 49, 784–797 (2021). PMID: 33653171.

18. Kuklyte, J. et al. Evaluation of the use of single- and multi-magnification convolutional neural networks for the determination and quantitation of lesions in nonclinical pathology studies. Toxicologic Pathology 49, 815–842 (2021).

19. Baek, E. B. et al. Artificial Intelligence-Assisted image analysis of Acetaminophen-Induced acute hepatic injury in Sprague-Dawley rats. Diagnostics (Basel) 12 (2022).

20. Bertani, V., Blanck, O., Guignard, D., Schorsch, F. & Pischon, H. Artificial intelligence in toxicological pathology: Quantitative evaluation of compound-induced follicular cell hypertrophy in rat thyroid gland using deep learning models. Toxicologic Pathology 50, 23–34 (2022). PMID: 34670459.

21. Mehrvar, S. et al. Deep learning approaches and applications in toxicologic histopathology: Current status and future perspectives. Journal of Pathology Informatics 12, 42 (2021).

22. Shimazaki, T. et al. Deep learning-based image-analysis algorithm for classification and quantification of multiple histopathological lesions in rat liver. Journal of Toxicologic Pathology 35, 135–147 (2022).

23. Gámez Serna, C., et al. Mmo-net (multi-magnification organ network): A use case for organ identification using multiple magnifications in preclinical pathology studies. Journal of Pathology Informatics 13, 100126 (2022).

24. Pischon, H. et al. Artificial intelligence in toxicologic pathology: Quantitative evaluation of compound-induced hepatocellular hypertrophy in rats. Toxicologic Pathology 49, 928–937 (2021).

25. Hwang, J.-H. et al. A comparative study on the implementation of deep learning algorithms for detection of hepatic necrosis in toxicity studies. Toxicological Research 39, 399–408 (2023).

26. Klambauer, G., Clevert, D.-A., Shah, I., Benfenati, E. & Tetko, I. V. Introduction to the special issue: Ai meets toxicology. Chemical Research in Toxicology 36, 1163–1167 (2023). PMID: 37599584.

27. Azizi, S. et al. Robust and data-efficient generalization of self-supervised machine learning for diagnostic imaging. Nature Biomedical Engineering 1–24 (2023).

28. Huang, Z., Bianchi, F., Yuksekgonul, M., Montine, T. & Zou, J. A visual–language foundation model for pathology image analysis using medical twitter. Nature Medicine 29, 1–10 (2023).

29. Chen, R. J. et al. Towards a general-purpose foundation model for computational pathology. Nature Medicine 1–13 (2024).

30. Xu, H. et al. A whole-slide foundation model for digital pathology from real-world data. Nature (2024).

31. Zhou, J., et al. ibot: Image bert pre-training with online tokenizer. International Conference on Learning Representations (ICLR) (2022).

32. Oquab, M., et al. DINOv2: Learning robust visual features without supervision. Transactions on Machine Learning Research (2024).

33. Caron, M. et al. Emerging properties in self-supervised vision transformers. In 2021 IEEE/CVF International Conference on Computer Vision (ICCV), 9630–9640 (2021).

34. Igarashi, Y. et al. Open TG-GATEs: a large-scale toxicogenomics database. Nucleic Acids Research 43, D921–D927 (2014).

35. Ilse, M., Tomczak, J. & Welling, M. Attention-based deep multiple instance learning. In Proceedings of the 35th International Conference on Machine Learning, 2132–2141 (2018).

36. Lu, M. Y. et al. Data efficient and weakly supervised computational pathology on whole slide images. Nature Biomedical Engineering (2020).

37. He, K., Zhang, X., Ren, S. & Sun, J. Deep residual learning for image recognition. In Proceedings of the IEEE conference on computer vision and pattern recognition, 770–778 (2016).

38. Shao, Z. et al. Transmil: Transformer based correlated multiple instance learning for whole slide image classification. Advances in Neural Information Processing Systems 34 (2021).

39. Wang, X. et al. Transpath: Transformer-based self-supervised learning for histopathological image classification. In International Conference on Medical Image Computing and Computer-Assisted Intervention, 186–195 (Springer, 2021).

40. Thoolen, B. et al. Proliferative and Nonproliferative Lesions of the Rat and Mouse Hepatobiliary System. Toxicologic Pathology 38, 5S–81S (2010).

41. Waters, M. D. & Fostel, J. M. Toxicogenomics and systems toxicology: aims and prospects. Nat. Rev. Genet. 5, 936–948 (2004).

42. Nyström-Persson, J., Natsume-Kitatani, Y., Igarashi, Y., Satoh, D. & Mizuguchi, K. Interactive Toxicogenomics: Gene set discovery, clustering and analysis in Toxygates. Scientific Reports 7, 1–10 (2017).

43. Hoeng, J., et al. *Hayes’ Principles and Methods of Toxicology*, chap. Toxicopanomics: Applications of Genomics, Transcriptomics, Proteomics, and Lipidomics in Predictive Mechanistic Toxicology (2023).

44. Varga, O. E., Hansen, A. K., Sandøe, P. & Olsson, I. A. S. Validating Animal Models for Preclinical Research: A Scientific and Ethical Discussion. Alternatives to Laboratory Animals 38, 245–248 (2010).

45. Moulin, P. et al. Imi—bigpicture: A central repository for digital pathology. Journal of Toxicologic Pathology (2021).

46. Vaswani, A. et al. Attention is all you need. Advances in neural information processing systems 30 (2017).

47. Dosovitskiy, A., et al. An image is worth 16x16 words: Transformers for image recognition at scale. In International Conference on Learning Representations (2021).

48. Devlin, J., Chang, M.-W., Lee, K. & Toutanova, K. Bert: Pre-training of deep bidirectional transformers for language understanding. Proceedings of the 2019 Conference of the North American Chapter of the Association for Computational Linguistics: Human Language Technologies, Volume 1 (Long and Short Papers) (2018).

49. Bao, H., Dong, L., Piao, S. & Wei, F. BEit: BERT pre-training of image transformers. In International Conference on Learning Representations (2022).

50. Garrido, Q., Balestriero, R., Najman, L. & Lecun, Y. Rankme: Assessing the downstream performance of pretrained self-supervised representations by their rank. In International conference on machine learning (2022).

51. Bankhead, P., et al. Qupath: Open source software for digital pathology image analysis. Scientific Reports 7 (2017).

52. Deng, J. et al. Imagenet: A large-scale hierarchical image database. In 2009 *IEEE conference on computer vision and pattern recognition*, 248–255 (Ieee, 2009).

53. Lu, M. Y. et al. Data-efficient and weakly supervised computational pathology on wholeslide images. Nature biomedical engineering 5, 555–570 (2021).

54. Chen, R. J. et al. Multimodal co-attention transformer for survival prediction in gigapixel whole slide images. In Proceedings of the IEEE/CVF International Conference on Computer Vision, 4015–4025 (2021).

55. Wang, X. et al. Transformer-based unsupervised contrastive learning for histopathological image classification. Medical Image Analysis 81, 102559 (2022).

56. Liu, Z. et al. Swin transformer: Hierarchical vision transformer using shifted windows. In Proceedings of the IEEE/CVF International Conference on Computer Vision, 10012–10022 (2021).

57. Kim, Y. J. et al. Paip 2019: Liver cancer segmentation challenge. Medical Image Analysis 67, 101854 (2021).

58. Chen, X*., Xie, S. & He, K*. An empirical study of training self-supervised vision transformers. In 2021 IEEE/CVF International Conference on Computer Vision (ICCV) (2021).

59. Zaheer, M. et al. Deep sets. Advances in Neural Information Processing Systems (2017).

60. Javed, S. A., et al. Additive mil: Intrinsically interpretable multiple instance learning for pathology. In Advances in Neural Information Processing Systems (NeurIPS) (2022).

61. Liu, L., et al. On the variance of the adaptive learning rate and beyond. In International Conference on Learning Representations (2020).

62. Snell, J., Swersky, K. & Zemel, R. Prototypical networks for few-shot learning. Advances in neural information processing systems 30 (2017).

63. Wang, Y., Chao, W.-L., Weinberger, K. Q. & Van Der Maaten, L. Simpleshot: Revisiting nearest-neighbor classification for few-shot learning. arXiv preprint arXiv:1911.04623 (2019).

64. Tian, Y., Wang, Y., Krishnan, D., Tenenbaum, J. B. & Isola, P. Rethinking few-shot image classification: a good embedding is all you need? In Computer Vision–ECCV 2020: 16th European Conference, Glasgow, UK, August 23–28, 2020, Proceedings, Part XIV 16, 266–282 (Springer, 2020).

